# Rapid depletion and super-resolution microscopy reveal an unexpected role of the nuclear-speckle protein SRSF5 in paraspeckle assembly and dynamics during cellular stress

**DOI:** 10.1101/2024.08.11.607506

**Authors:** Benjamin Arnold, Laurell Kessler, Ellen Kazumi Okuda, Ricarda J. Riegger, Maria Clara Hernández Cañás, Ewelina Zebrowska, Cem Bakisoglu, Helder Y. Nagasse, David Stanek, Dorothee Dormann, Kathi Zarnack, Mike Heilemann, Michaela Müller-McNicoll

**Author notes:** These authors contributed equally and appear in alphabetical order (Benjamin Arnold, Laurell Kessler, Ellen Kazumi Okuda).

## Abstract

Nuclear speckles (NS) and paraspeckles (PS) are adjacent condensates with distinct protein composition, with serine-arginine-rich splicing factors (SRSFs) concentrated in NS. Surprisingly, we find that SRSF5 is present in both. Combining super-resolution imaging, proximity proteomics and iCLIP, we show that SRSF5 binds with PS core proteins to the PS-scaffold RNA *NEAT1* and locates between PS spheres. Acute SRSF5 depletion results in reduced PS with differently packaged *NEAT1*. Under stress, SRSF5’s association with PS increases, and without SRSF5, PS cluster assembly is impaired. Interfering with binding to purine-rich RNAs even causes PS-NS fusion. In an intriguing over-compensation, longer SRSF5 depletion reduces TDP-43 levels via premature polyadenylation, leading to *NEAT1* isoform switching and more PS. We propose that SRSF5 forms a stress-specific PS shell and acts as a glue for PS clusters. Additionally, we uncover SRSF5 as a novel regulator of TDP-43 and demonstrate how acute depletion distinguishes direct from compensatory effects.

**Highlights:**

- NS protein SRSF5 associates with PS shells and enriches between PS spheres
- SRSF5 binds the PS-scaffold RNA *NEAT1* and ensures proper *NEAT1* packaging
- SRSF5 association with PS increases under stress and promotes cluster formation
- Interfering with binding to purine-rich RNAs causes the fusion of PS and NS
- Acute SRSF5 depletion reveals compensatory effects on PS assembly and dynamics
- SRSF5 regulates TDP-43 levels via premature polyadenylation

## Introduction

The nucleus of mammalian cells is compartmentalized into distinct phase-separated membrane-less organelles (MLOs) to spatially organize concurring synthesis and processing pathways and to regulate gene expression in response to stress ^1–3^. One of the main hubs for the organization of the cellular stress response in space and time are paraspeckles (PS) ^4^. PS are scaffolded by the architectural RNA *NEAT1* and contain 80 different RNA-binding proteins (RBPs) ^5–8^, of which eight are essential for PS formation (DAZAP1, FUS, HNRNPH3, HNRNPK, NONO, RBM14, SFPQ, and SMARCA4) ^9–11^. Most PS proteins (PSPs) contain intrinsically disordered regions (IDRs) or prion-like domains, which are essential for PS assembly and promote phase separation ^12,13^. The precise composition of PS is not known, but it is thought to be dynamic and stress-dependent, to be able to sequester specific RNAs and proteins in response to different types of cellular stress ^14^.

PS assemble co-transcriptionally by PSPs binding to nascent *NEAT1* transcripts in a coordinated manner, which forces *NEAT1* to adopt a V-shaped structure and keeps monomeric *NEAT1* ribonucleoprotein particles (RNPs) in close proximity. Approximately 50 *NEAT1* RNPs then assemble into a mature phase-separated PS with an intricate core-shell structure and a fixed diameter of 360 nm ^15–18^. The 5’ and 3’ends of *NEAT1* are facing the shell, and the middle region is hidden in the PS core, while the PSP FUS acts as a separator of individual chains ^17^.

*NEAT1* exists in two major isoforms that share the same 5’end region, but only the long isoform (*NEAT1_2*; 22 kb) scaffolds PS and their numbers strictly correlate. While the short *NEAT1* isoform (*NEAT1_1;* 3.7 kb) is canonically polyadenylated, the long isoform only forms when polyadenylation of *NEAT1_1* is prevented. *NEAT1_2* harbors a tRNA-like module at its 3’end, which is cleaved off by RNase P. The resulting 3’end is stabilized through a U–A°U triple-helix structure ^8,19^. Several RBPs are known to regulate the ratio between both isoforms, such as HNRNPK that competes with cleavage factor CPSF6 for the binding of NUDT21 and thereby prevents *NEAT1_1* formation ^11^. Conversely, TDP-43 and the Integrator subunits INTS10 and INTS11 promote *NEAT1_1* formation by enhancing its polyadenylation ^20,21^.

During stress, PS increase in size and number and form larger clusters through increased *NEAT1* transcription and enhanced polyadenylation site (PAS) read-through ^1,4^. The individual PS grow in only one dimension and form cylinder-shaped structures, explainable by a co-polymer micelle model ^22,23^, but the architecture of larger PS clusters is not understood.

PS are often found in close proximity to nuclear speckles (NS), which are nuclear MLOs that sequester splicing factors and regulate co-transcriptional mRNA processing ^6,24^. Of note, the RBPs that reside in PS and NS are fundamentally different. NS contain RBPs that harbor domains rich in arginine and serine residues (RS domains) – including serine-arginine-rich splicing factors (SRSFs) – that form a tight meshwork and assemble the NS core ^25^. The NS phase border is populated by active spliceosomes and poly(A)+ RNAs ^26,27^. In contrast, repressory splicing factors – including heterogeneous nuclear ribonucleoproteins (hnRNPs) – are expelled from NS but are enriched in PS. Thus, the types of RBPs and their interactions provide both MLOs with very different physico-chemical properties and might prevent their fusion. Indeed, a recent study demonstrated that RBPs that localize to the shell of PS, e.g. SFPQ are important for the separation of both MLOs ^28^, but it is less clear how PS and NS are held in close proximity. However, one shared component of both MLOs might be purine-rich RNAs. Purine-rich RNAs are enriched in NS and bound by SR proteins that localize to the core of NS ^29,30^. But AG-rich RNAs were also found enriched in PS and distributed along the shell of PS spheroids ^17^.

We have recently shown that during hypoxia stress, NS partially dissolve and release SRSFs, while at the same time PS increase in size and number ^31^. This raises the possibility that NS and PS do crosstalk and exchange components during the stress response. To investigate a potential crosstalk, we focused on SRSF5, which is a known NS core component and binds to purine-rich RNAs, but was also recently found in PS by mass spectrometry ^32^ and shown to bind to *NEAT1* ^33,34^. The role of SRSF5 in PS biology or crosstalk with NS is unknown.

Here, we show that SRSF5 is critical to regulate PS assembly by both direct and indirect mechanisms. We demonstrate that SRSF5 localizes to PS shells and promotes proper *NEAT1_2* packaging and PS cluster formation during certain stress conditions. SRSF5 also regulates PS indirectly by regulating the levels of TDP-43 via premature polyadenylation, which drives a switch in *NEAT1* isoforms in response to prolonged SRSF5 depletion.

## Results

### SRSF5 associates with nuclear speckles and paraspeckles

A recent study suggested that the NS-resident protein SRSF5 also locates to PS ^32^. To confirm this association we first examined the subnuclear localization of endogenously tagged SRSF5-GFP in HeLa cells ^35^. NS were detected by immunofluorescence for the NS marker SRRM2 and PS by RNA fluorescence *in situ* hybridization (RNA-FISH) with probes hybridizing to the middle region of *NEAT1_2*. Confocal microscopy revealed that SRSF5 localized primarily to NS (63%) but also to the nucleoplasm (37%) (**Figure 1A**). While 36% of PS signal was far away from SRSF5 or NS (**Figure 1B**), 47% of SRSF5 signal was found in close proximity and 17% overlapped with PS, suggesting that SRSF5 often lies between neighboring NS and PS.

**Figure 1.**
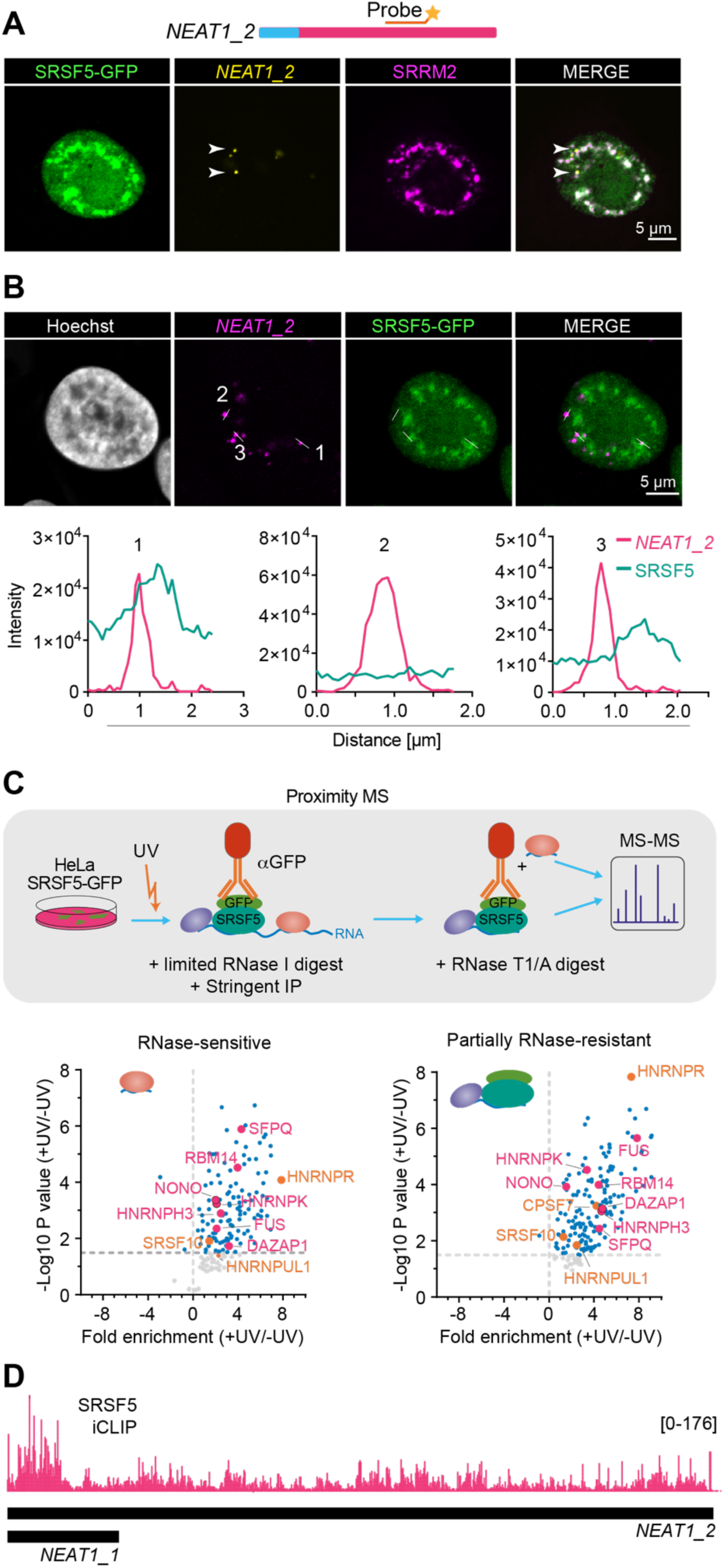
SRSF5 associates with paraspeckles and nuclear speckles. **A)** Confocal images of HeLa SRSF5-GFP cells showing that SRSF5 localizes to NS, labelled with an anti-SRRM2 antibody. PS were labelled by RNA-FISH targeting the middle region of *NEAT1_2*. **B) Top:** Confocal images of HeLa SRSF5-GFP cells showing that SRSF5 signal overlaps with PS. PS were labelled by RNA-FISH targeting the middle region of *NEAT1_2*. Nuclei were stained with Hoechst. **Bottom:** Example line scans show partial overlap (1), no co-localization (2) and close proximity (3) of SRSF5-GFP and *NEAT1_2*. **C) Top:** Scheme of the proximity MS workflow that identified all proteins that bind in close proximity to SRSF5 to the same short fragment (150 nt) of RNA^36^. **Bottom:** RNase-sensitive (left) and partial RNase-resistant (right) SRSF5 pull-downs reveal that SRSF5 binds in proximity to most known PSPs. Essential PSPs - pink, important PSPs - orange. **D)** Browser shot showing the preferential binding of SRSF5 at the 5’end of the *NEAT1_2* transcript determined by iCLIP2^35^.

We next re-analyzed our previous proximity mass spectrometry (MS) dataset ^36^ that identified all proteins binding together with SRSF5 on short RNA fragments, which is based on UV-crosslinking, titrated RNase digestion and SRSF5 pull-down. This confirmed that most known PSPs including FUS, HNRNPK, NONO, RBM14, and SFPQ were strongly enriched in UV-crosslinked SRSF5 pull-downs (compared to -UV) both in RNase-sensitive (150 nt) and partially RNase-resistant (same complex) samples (**Figure 1C**). To test whether these interactions occurred within PS, we inspected a recent iCLIP2 dataset performed with the same cell line ^35^ and found that SRSF5 binds strongly to the PS-scaffolding RNA *NEAT1_2* and preferentially to its 5’end (**Figure 1D**). This would place SRSF5 to the shell region of PS spheres. Collectively, these data suggest that SRSF5 is associated not only with NS, but also with PS in steady-state conditions.

### Acute depletion of SRSF5 decreases paraspeckle number and size

To address whether SRSF5 plays a role in PS biology, we employed hGRAD, our recently developed doxycycline-inducible system for rapid degradation of MLO-associated RBPs tagged with GFP ^35^. Treatment of HeLa hGRAD cell lines expressing endogenously GFP-tagged SRSF5 with doxycycline (DOX; 1 µg/mL) induced hGRAD-mediated rapid degradation of the protein within 2 h (**Figure 2A, S1**). We found that acute SRSF5 depletion (∼8 h) had no effects on the numbers and size of NS (**Figure 2G**). Interestingly, however, we observed a dramatic decrease in the number and size of PS. In fact, PS virtually disappeared (**Figures 2A-C**). Moreover, acute depletion of SRSF5 caused a slight but significant decrease in the distance between NS and remaining PS (**Figure 2G, 2H**). The observed effects were specific to SRSF5, as the depletion of the NS core protein SRRM2 did not affect PS (**Figure 2E**), supporting that PS disappearance was neither due to the induction of the hGRAD system per se or nor due to depletion of an NS core protein (SRRM2). In contrast, the acute removal of the PS core protein NONO disassembled PS as expected, but small evenly distributed *NEAT1_2* foci remained visible in these cells (**Figure 2F**). Altogether, this suggests that SRSF5 is important for PS assembly or stability and influences the distance to NS and the crosstalk between the two MLOs (**Figure 2D**).

**Figure 2.**
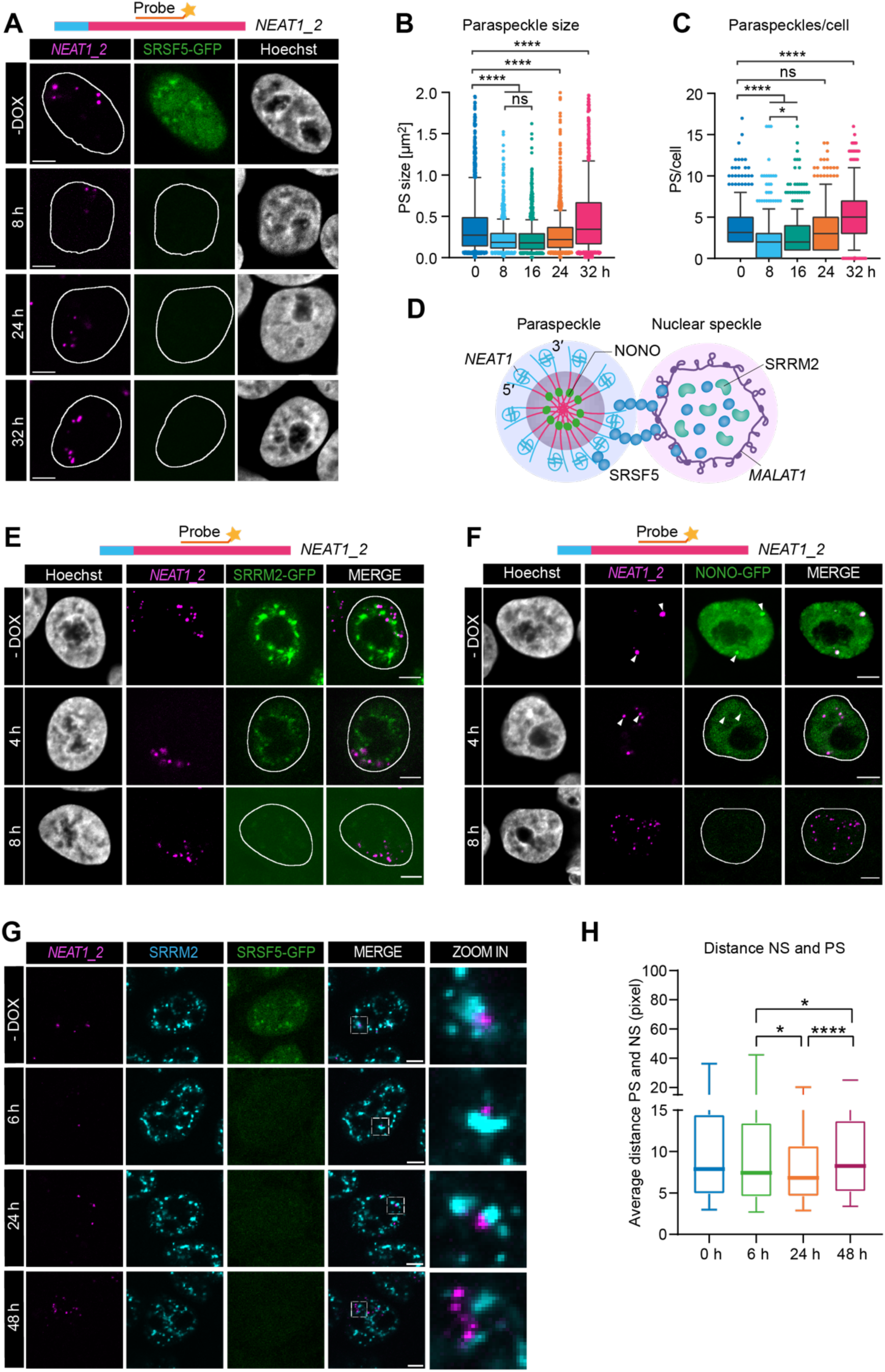
Acute depletion of SRSF5 decreases paraspeckle number and size. **A)** Scheme of the architecture of PS and NS with indicated marker RNAs *NEAT1* and *MALAT1* and RBPs NONO, SRRM2 and SRSF5. **B)** Confocal images showing a time course of SRSF5-GFP depletion (8, 24 and 32 h) using hGRAD. **C)** Quantification of PS size from n = 319-375 cells and **D)** PS number per cell from n = 319-375 cells upon hGRAD-mediated depletion of SRSF5-GFP. Statistics: two-tailed Mann-Whitney test. P-value: ≤ 0.05 (*), < 0.0001 (****) **E-F)** Time course of SRRM2-GFP (E) and NONO-GFP (F) depletion (-DOX, 4 and 8 h) using hGRAD. **G)** Time course of SRSF5-GFP depletion (6, 24 and 48 h) using hGRAD. SRRM2 was immuno-stained with an anti-SRRM2 antibody. **H)** Quantifications of the average distances between PS and NS from n = 549-977 PS. Statistics: two-tailed Mann-Whitney test. All PS were labelled by RNA-FISH targeting the middle region of *NEAT1_2.* Scale bars - 5 µm.

### Paraspeckles are smaller and differently packaged in the absence of SRSF5

To explore SRSF5’s association with PS with higher resolution, we visualized individual PS in HeLa cells using DNA points accumulation for imaging in nanoscale topography (DNA-PAINT) ^37,38^ with RNA-FISH probes that hybridize to the 5’end of *NEAT1_2* and direct Stochastic Optical Reconstruction Microscopy (dSTORM). At higher resolution of dSTORM, PS became visible also during acute SRSF5 depletion (6 h), but these were significantly smaller in diameter (317 nm) than control PS in untreated cells (362 nm; **Figure 3A**). Moreover, we found mainly spheres but no larger PS clusters (**Figure 3B**), together explaining why PS escaped detection by standard confocal microscopy.

**Figure 3.**
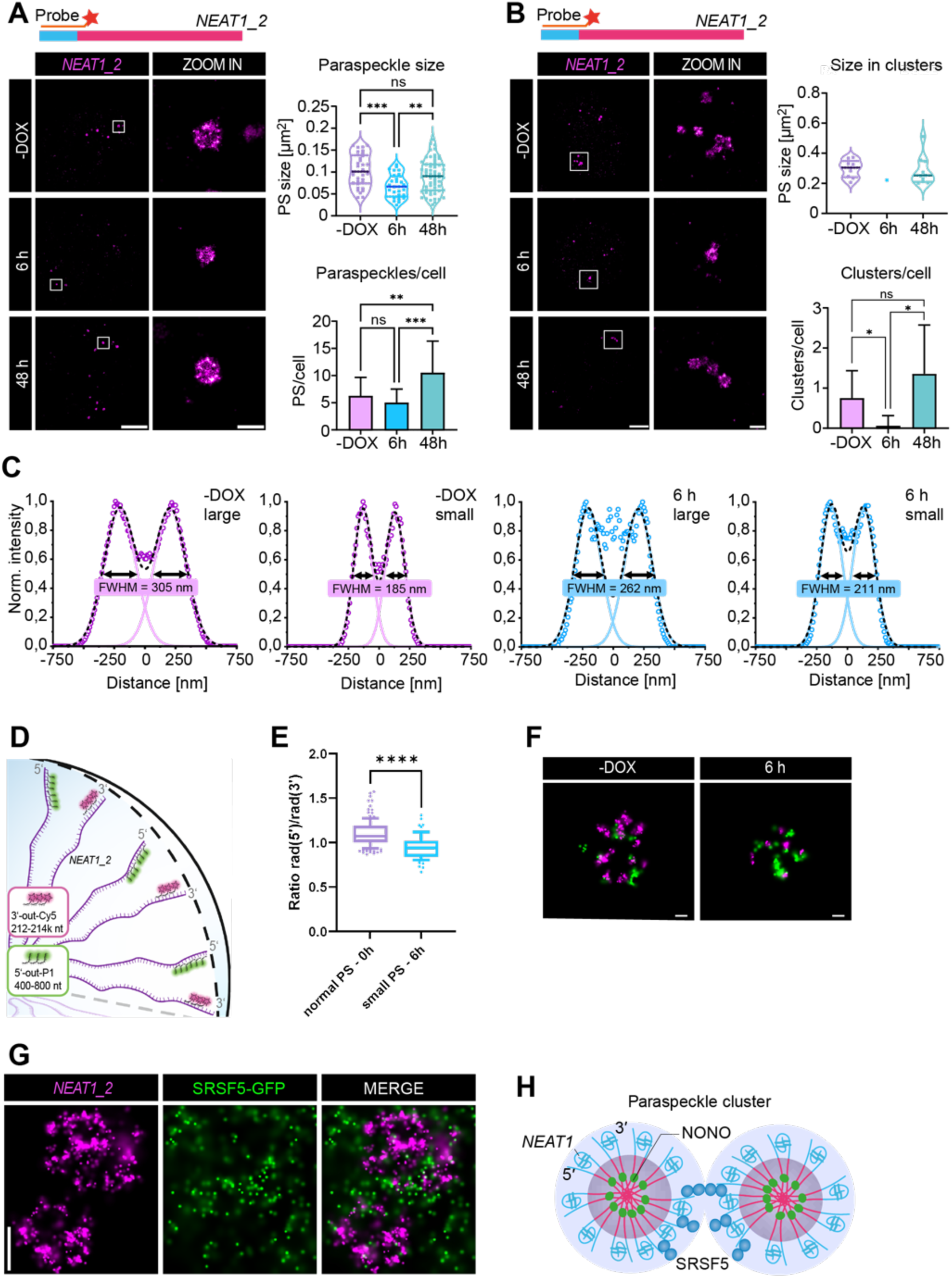
Paraspeckles are smaller and differently packaged in the absence of SRSF5. **A-C)** Direct Stochastic Optical Reconstruction Microscopy (dSTORM) imaging PS after 6 h and 48 h SRSF5-GFP depletion using hGRAD. PS were labelled by DNA-PAINT RNA-FISH using probes targeting the 5’end of *NEAT1_2.* **A)** Quantification of PS size and PS number in PS spheres from n = 8 cells. Scale bars - 5 µm and 500 nm. **B)** Quantification of PS that are part of clusters and number of clusters per cell from n = 7-9 cells. Scale bars – 5 µm and 500 nm. **C)** Distance distribution of normalized 5’end signal from the center of large and small PS comparing control (0 h) and SRSF5-depleted (6 h, 1 µg/mL DOX) cells. FWHM - full width at half maximum. n=3-8 PS **D)** Multicolor DNA-PAINT RNA-FISH was used to label the 5’ and 3’end of *NEAT1*. Positions of where the probes bind to the *NEAT1_2* RNA are indicated. **E)** Quantification of the ratios between 5’ and 3’ probes. **F)** dSTORM image of representative PS spheres at 0 h at 6 h SRSF5 depletion. **G)** DNA-PAINT RNA-FISH using probes at the 5’end of *NEAT1_2* was coupled with dSTORM imaging for SRSF5-GFP using GFP nanobodies. Scale bars - 200 nm. **H)** Current model of *NEAT1_2* folding and packaging. SRSF5 binding to the 5’end of *NEAT1_2* locates to the shell of PS where it could mediate the interactions between different PS spheres to assemble larger clusters.

The diameter of PS is supposed to be invariant (360 nm) due to the fixed length and packaging of *NEAT1_2* molecules inside PS, whereby the 5’ and 3’end of *NEAT1_2* localize to the shell ^16,28^ (**Figure 2A, 3D**). Thus, smaller PS suggest that in the acute absence of SRSF5, *NEAT1_2* RNAs are differently packaged. To test this, we compared the distance distribution of the 5’end signal of *NEAT1_2* in control (0 h) and 6 h SRSF5-depleted cells (**Figure 3C**). Since smaller spheres were also found in control conditions, although much less frequent, we separated PS into large and small objects and measured their width (full width at half maximum (FWHM)). In control conditions large objects showed a FWHM of 305 nm and small objects of 185 nm. After 6 h SRSF5 depletion, the FWHM decreased strongly for large objects from 305 to 262 nm, indicating that PS become smaller and that the 5’end is more compacted. Such a decrease was not seen for already small particles (185 to 211 nm). Interestingly, while in control cells 5’end signal density was completely depleted in the center part of PS (core), in SRSF5-depleted cells (6 h), both, large and small objects showed 5’end signal of *NEAT1_2* in the center of PS. This suggests that the shell localization of *NEAT1_2* is partially impaired upon SRSF5 depletion and *NEAT1_2* 5’ends partially mis-localize to the PS core (**Figure 3C**).

To further evaluate differences in the structural organization of PS, we labeled the 5’ and 3’end of *NEAT1_2* simultaneously, recorded super-resolution Stimulated Emission Depletion (STED) data ^37,39–41^ and performed a single-particle analysis (see Methods; **Figure 3D, S2A**). We observed a statistically significant decrease in the ratio of 3’ to 5’signal for SRSF5-depleted PS (6 h, **Figure 3E**), which suggests that the 5’end was indeed more internalized (with respect to the 3’ label). Example PS imaged with dSTORM also suggested that PS were smaller and the 5’end were more internalized compared to control PS (**Figure 3F**), which was not observed using two neighboring probes at the 3’end of *NEAT1_2* (**Figure S2B**).

Super-resolution dSTORM imaging of SRSF5-GFP and *NEAT1_2* revealed that SRSF5 decorated the outside of PS but was particularly enriched between PS spheres (**Figure 3G, S2C**). Together with the preferred binding of SRSF5 to the 5’end of *NEAT1_2*, our data suggest that SRSF5 is required for the shell localization of the 5’end and that PS packaging occurs correctly, as suggested for other shell proteins ^9^. It may further allow the assembly of larger PS clusters through SRSF5-mediated interactions between shell regions (**Figure 3H**). SRSF5 depletion alters the *NEAT1_2* packaging and causes smaller PS that can no longer assemble into larger clusters.

### Long-term depletion or SRSF5 KO trigger a PS over-compensation mechanism

While our data using acute depletion suggested that SRSF5 plays a role for the correct assembly of PS, longer SRSF5 depletion times (32-48 h) revealed a surprising recovery of PS. Their sizes and numbers as well as the distance to NS increased at longer depletion times (32 h) compared to control cells (0 h; **Figures 2B-D, 2G, 2H**). This recovery was confirmed by dSTORM super-resolution imaging using probes at the 5’end of *NEAT1_2*. After 48 h of SRSF5 depletion, PS were more numerous than in control cells (0 h), they recovered their original diameter (360 nm) and more often assembled in large PS clusters (**Figure 3A, 3B**).

To investigate this phenomenon further, we generated a HeLa SRSF5 knockout (KO) cell line using CRISPR/Cas9 (**Figure S3A-C**). Confocal and dSTORM super-resolution imaging revealed that similar to long-term depletion by hGRAD, KO cells showed much more PS than WT cells (**Figure 4A-D**), they exhibited more PS clusters and elongated structures and PS spheres had a normal diameter of 360 nm (**Figure 4D, 4E**). Moreover, the distance to NS increased compared to WT cells (**Figure S3D**). Altogether these data suggest that the effects of acute and long-term depletion of SRSF5 differ and the latter may trigger a compensation mechanism to restore PS assembly.

**Figure 4.**
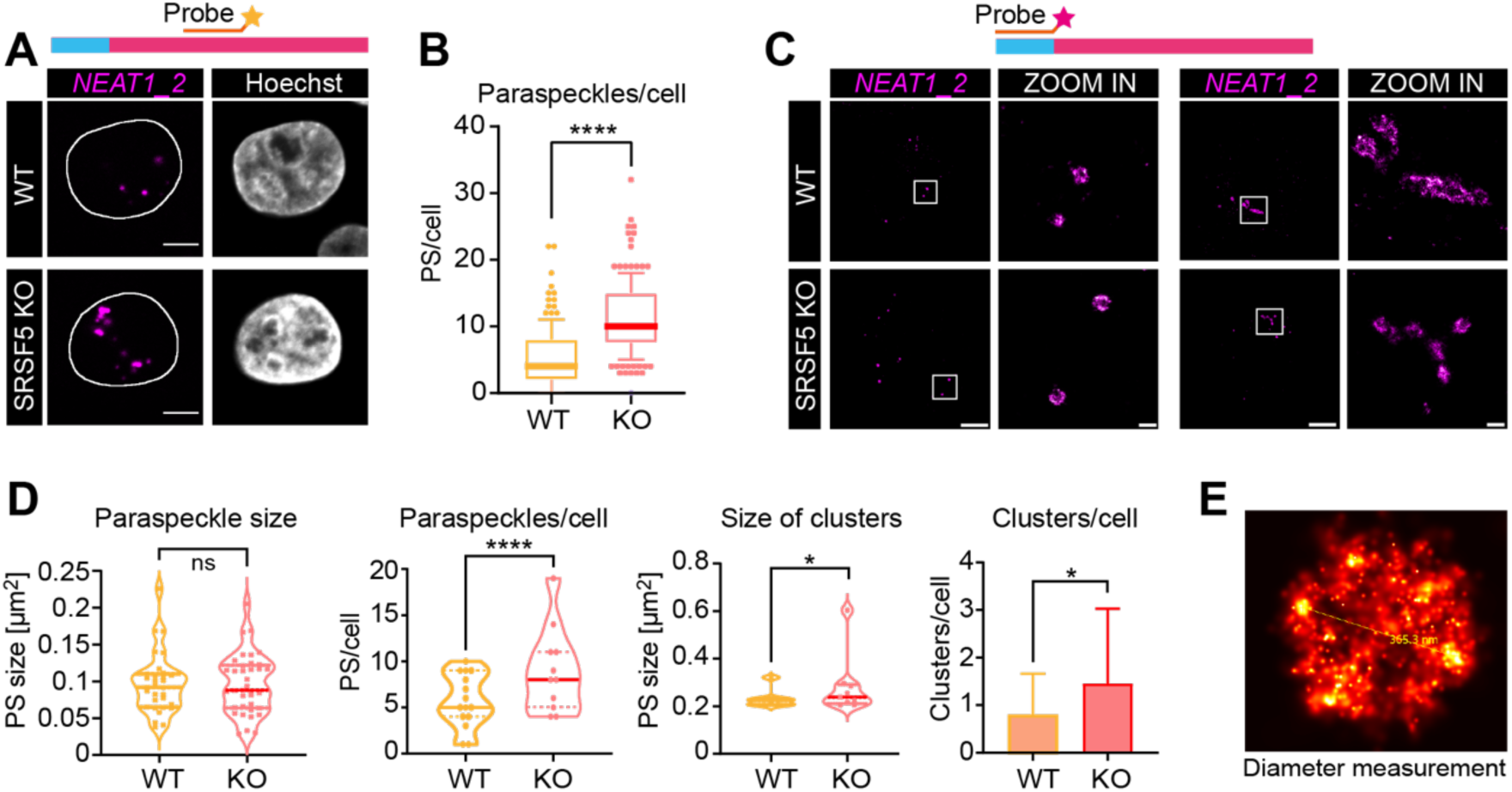
SRSF5 KO triggers a PS over-compensation mechanism. **A)** Confocal imaging comparing PS number in HeLa WT and SRSF5 KO cells. PS were labelled by RNA-FISH using probes hybridizing to the middle region of *NEAT1_2.* Scale bars - 5 µm. **B)** Quantification of PS number per cell from n = 300 cells. T-test and stdev are indicated. **** p < 0.0001. **C)** dSTORM imaging of WT and SRSF5 KO cells. PSs were labelled by DNA-PAINT RNA-FISH using probes hybridizing to the 5’end of *NEAT1_2.* Scale bars - 5 µm and 500 nm. **D)** Quantification of the size of PS spheres, the number of PS spheres per cell, the size of PS clusters and the number of PS clusters per cell from n = 9 cells. T-test and stdev are indicated. * p < 0.05; ** p < 0.01; *** p < 0.001; **** p < 0.0001. **E)** Example micrograph from one PS sphere with a diameter of 365 nm using probes hybridizing to the 5’end of *NEAT1_2*.

### Acute SRSF5 depletion destabilizes NEAT1_2 transcripts

To understand the nature of this over-compensation, we analyzed our previously generated Nascent-seq dataset where we had combined acute SRSF5 depletion by hGRAD with nascent RNA sequencing ^35,42^. For this dataset, HeLa hGRAD cells had been induced by DOX addition and after different time points of SRSF5 depletion (2 h - T2; 8 h - T8; 16 h - T16) they were treated with 400 µM 4sU for one hour. Only RNAs that were transcribed after SRSF5 depletion are labelled and sequencing allows to follow their expression dynamics and processing changes over time. Uninduced cells (T0) served as control. To investigate changes upon long-term SRSF5 depletion, we additionally prepared a Nascent-seq dataset for HeLa SRSF5 KO and WT cells (**Figure 5A**). Differential expression analysis using DESeq2 ^43^ revealed very rapid and dynamic changes in the expression of hundreds of transcripts upon removal of SRSF5 (**Figure 5B**). One of the transcripts with such a dynamic behavior was *NEAT1*. *NEAT1* is down-regulated at T2 and T8, recovered at T16 and increased by two-fold in SRSF5 KO cells (**Figure 5C**). These changes in *NEAT1* levels precisely mirror the observed changes in PS over time (**Figure 2A**). The initial decrease of *NEAT1* levels and later increase at 32 h depletion and in SRSF5 KO cells was also seen by RT-qPCR performed with total RNA (**Figure 5D, 5E**). This shows that not only the newly transcribed *NEAT1_2* transcripts disappear upon acute depletion of SRSF5, but also older molecules decrease in abundance, pointing to a general loss of *NEAT1_2*. Thus, our data suggest that acute SRSF5 removal destabilizes *NEAT1_2* transcripts, but long-term SRSF5 depletion recovers its level.

**Figure 5.**
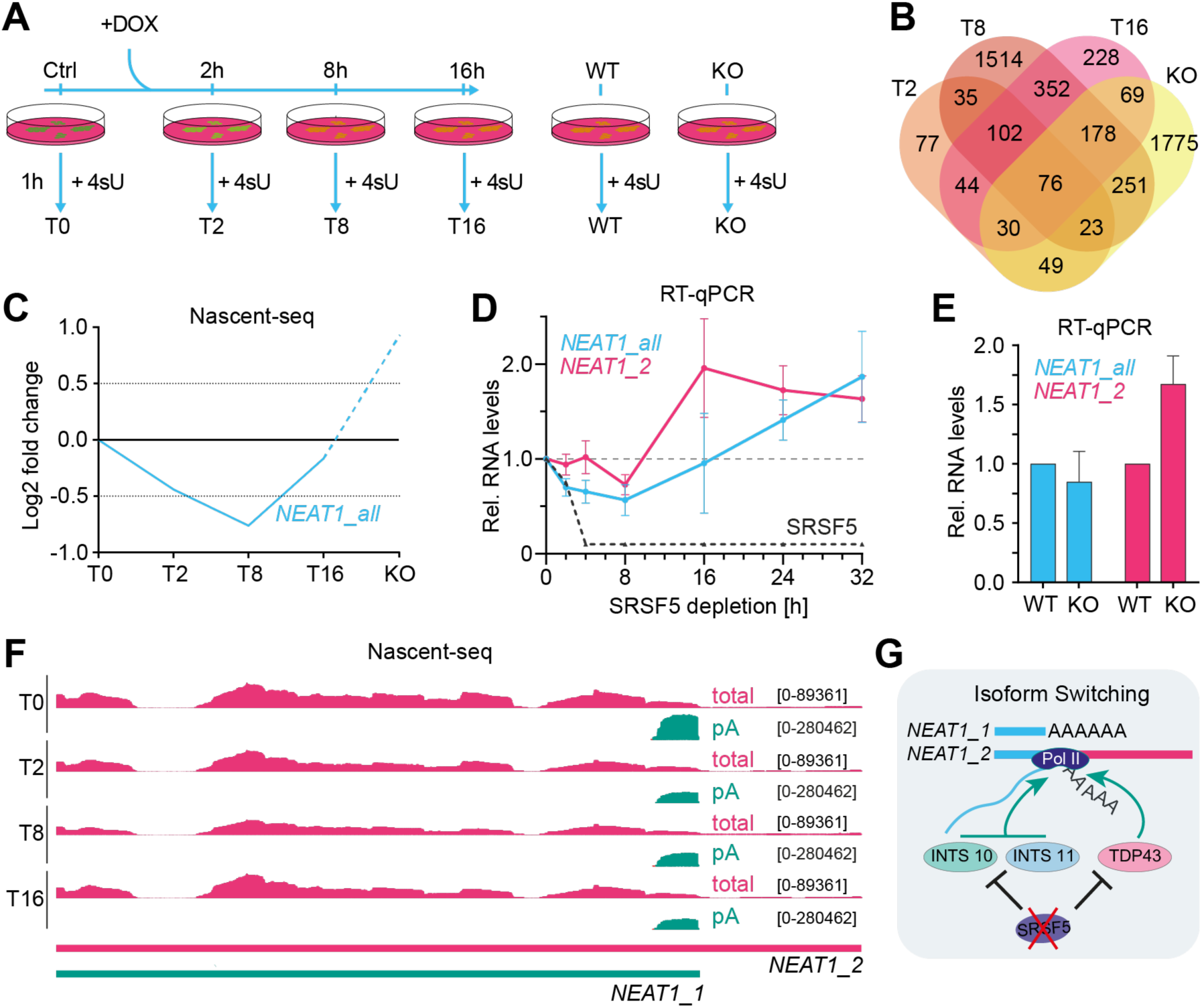
Over-compensation of paraspeckles is caused by an isoform switch to *NEAT1_2*. **A)** Scheme of the workflow combining a time course of hGRAD degradation as well as WT and SRSF5 KO with Nascent-seq. Cells were induced with DOX (1 µg/mL) for the indicated times, then treated for 1 h with 4sU (400 µM) to label all new transcripts followed by total RNA extraction, biotinylation, purification and conversion into a cDNA library for deep sequencing. **B)** Venn diagram of differentially expressed transcripts at T2, T8 and T16 compared to T0 and KO compared to WT. **C)** *NEAT1* shows dynamic transcript changes upon SRSF5 depletion. **D)** RT-qPCR confirms the dynamic *NEAT1* regulation after SRSF5 depletion with total RNA.**E)** RT-qPCR confirms the over-shooting of *NEAT1* levels in SRSF5 KO cells. RNA levels were normalized to U6 snRNA. Graphs show mean and SD of n=3 independent experiments. Statistics: two-tailed Mann-Whitney Test. **F)** Genome browser view showing total (pink) and polyA (green) Nascent-seq reads from the different time points of SRSF5 depletion mapping around the poly(A)site of *NEAT1_1*. The *NEAT1_2* transcript is truncated for better visibility. **G)** Model how SRSF5 affects *NEAT1* isoform switching.

### SRSF5 restricts NEAT1_2 production by maintaining high levels of TARDBP and INTS10

To address the compensatory mechanism that restores *NEAT1_2* levels after longer SRSF5 depletion times and KO, we inspected the ratios of the two *NEAT1* isoforms over time. The long *NEAT1_2* isoform, which scaffolds PS, is formed by polyadenylation signal read-through of the short *NEAT1_1* isoform^11,44^. Counting only reads with non-templated As (pA) that map to the *NEAT1* locus we found that the short polyadenylated *NEAT1_1* isoform was progressively decreasing after SRSF5 depletion (**Figure 5F, S4A**), while the long *NEAT1_2* isoform increased over time and remained elevated in SRSF5 KO cells (**Figure 5C, 5D**). Thus, later PS compensation may occur by PAS read-through and a progressive switch to the *NEAT1_2* isoform, which in turn can assemble more PS (**Figure 5G**). Several proteins are known to directly regulate the switch between the *NEAT1* isoforms by binding to the *NEAT1_1* polyadenylation site (PAS) and activating or inhibiting its usage. These include CPSF6, TDP-43, hnRNPK, INTS10 or INTS11 ^11,20,21^. However, SRSF5 may not be one of them, as the *NEAT1_1* PAS region is rather depleted of SRSF5 binding sites (**Figure S4B**).

PAS usage can also be regulated indirectly by altering the expression of CPA factors. For example, SRSF3 regulates the levels of CPSF6 by alternative splicing ^45,46^. To test this possibility, we evaluated expression changes of all known genes encoding *NEAT1* and PS regulators in the Nascent-seq dataset (**Figure S4C**). We found that the levels of *TARDBP*, which encodes for TDP-43, and *INTS11* encoding for one of the Integrator subunits, were progressively reduced upon SRSF5 depletion and in the SRSF5 KO, while the levels of *HNRNPK* and *CPSF6* did not change (**Figure 6A, S4C**). *INTS10* levels were also reduced at early time points of SRSF5 depletion, but recovered in SRSF5 KO cells (**Figure 6A**). We were able to validate the decrease of *INTS10* and *INTS11* mRNA levels until 30 h SRSF5 depletion by RT-qPCR using total RNA (**Figure 6B, 6C**), but not the decrease of *TARDBP,* suggesting that SRSF5 depletion affects only newly transcribed *TARDBP* mRNA and is masked by old transcripts. Nevertheless, *INTS11* and *TARDBP* mRNAs were decreased in the SRSF5 KO and the levels of TDP-43 and INTS10 proteins also progressively decreased in the absence of SRSF5 (**Figure 6D-F**). Since TDP-43 and INTS10 promote PAS usage and termination at the short *NEAT1_1* isoform^20,21^, their reduction may favor production of the long *NEAT1_2* isoform over time.

**Figure 6.**
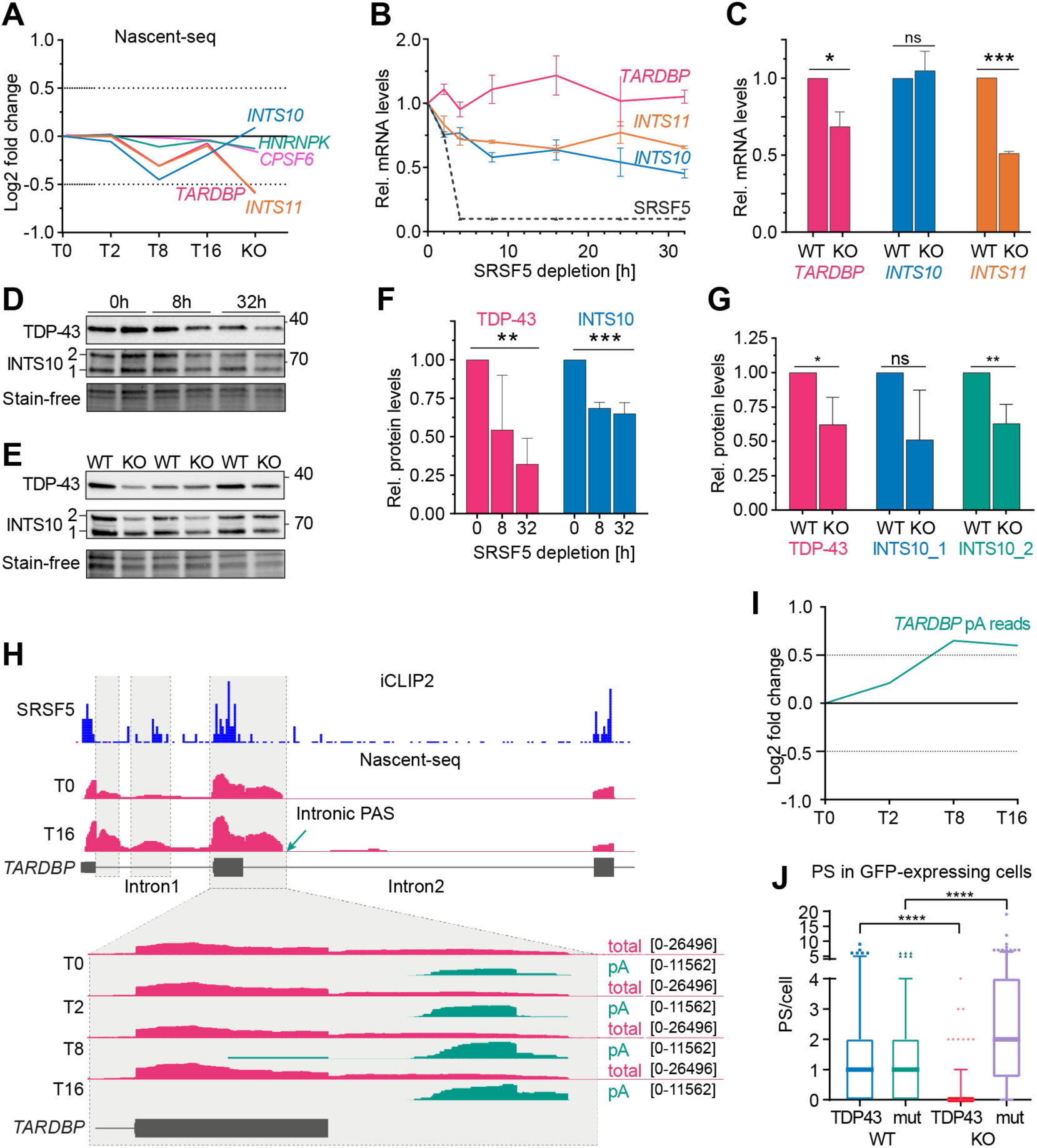
SRSF5 restricts *NEAT1_2* levels and PS assembly indirectly by regulating expression of TDP-43 and INTS10. **A)** Expression changes in genes encoding known PS regulators quantified from Nascent-seq data. **B)** RT-qPCR of a time course of SRSF5 depletion confirms the regulation of *TARDBP*, *INTS10* and *INTS11*. **C)** RT-qPCR confirms the regulation of *TARDBP*, *INTS10* and *INTS11* in SRSF5 KO cells. RNA levels were normalized to U6 snRNA. Graphs show mean and SD of n=3 independent experiments. Statistics: two-tailed Mann-Whitney Test. **D)** Representative Western blot of a time course of SRSF5 depletion showing the decrease in TDP-43 and INTS10 protein levels. Stain-free gels were used to control for equal loading. **E)** Representative Western blot of WT and SRSF5 KO cells showing the decrease in TDP-43 and INTS10 protein levels. Stain-free gels were used to control for equal loading. **F,G)** Quantification of TDP-43 and INTS10 protein levels of the Western blots partially shown in D and E from n=3 experiments. **H)** Genome browser view showing total (pink) and polyA (green) Nascent-seq reads mapping to the 5’end of the *TARDBP* transcript from the different time points of SRSF5 depletion. SRSF5 iCLIP2 crosslinks are shown in blue. **I)** Changes in the number of pA reads mapping to intron 2 of *TARDBP* quantified from Nascent-seq data. **J)** Overexpression of TDP-43 but not the cytoplasmic TDP43-mut decrease the number of paraspeckles in SRSF5 KO cells but not WT cells. Quantification from 3 independent replicates.

To investigate how SRSF5 depletion decreases the levels of *TARDBP*, *INTS10* and *INTS11* mRNA we analyzed the SRSF5 KO Nascent-seq data for alternative splicing using MAJIQ ^47^. This revealed 23 (T2), 159 (T8), 116 (T16) and 108 (KO) significant splicing events (**Figure S4D**). We did not detect any splicing changes in *INTS10* and *INTS11* transcripts, but intron 1 and 2 of *TARDBP* mRNA were progressively retained upon SRSF5 depletion. iCLIP2 data show that SRSF5 binds normally in exons flanking these retained introns (**Figure 6H**). Interestingly, retention of intron 2 activates an intronic PAS, which leads to early transcript termination. Quantification of polyadenylated reads (pA) mapping to the *TARDBP* locus revealed that this PAS was polyadenylated and its usage progressively increased (**Figure 6H, 6I**).

These data suggest that SRSF5 is a novel regulator of TDP-43 expression and promotes splicing of intron 1 and 2 of the *TARDBP* pre-mRNA. When SRSF5 is absent both introns are more retained, and an intronic PAS is used terminating transcription at this site. As a consequence, less TDP-43 protein is produced. This means that SRSF5 restricts *NEAT1_2* production and PS assembly indirectly by maintaining high levels of TDP-43 and INTS10.

To test whether PS over-compensation could be rescued in SRSF5 KO cells we overexpressed TDP-43 and a mutant version (TDP43-mut) that is retained in the cytoplasm fused to a GFP-tag in WT and KO cells using transient transfection (**Figure S5**). Quantification of PS number in cells expressing GFP revealed that TDP43 overexpression strongly suppressed PS number and size in SRSF5 KO cells but not in WT cells. The cytoplasmic TDP43 mutant version had no effect (**Figure 6J**), suggesting that PS over-compensation at long-term SRSF5 depletion is caused partially by a reduction in TDP-43 levels, which promotes *NEAT1_2* expression and assembly of more PS (**Figure 5G**).

### SRSF5 is required for PS hyper-assembly during stress

The main function of PS is to organize the cellular stress response in space and in time ^4^. *NEAT1* transcription is induced by many compounds or stress conditions resulting in large PS clusters that sequester PSPs and reduce their activities in transcription or splicing, which is important for the stress adaptation ^1,32^. Our data so far suggest that SRSF5 is a new shell protein that is required for PS cluster formation. However, PS cluster formation recovered after prolonged SRSF5 depletion, although those PS lacked SRSF5 in their shells. This suggests that PS with a different shell composition exist, which might respond differently to stress conditions as suggested previously ^4^.

To test whether these overcompensated PS are structurally different in WT and SRSF5 KO cells, we used dSTORM imaging with two different RNA-FISH probe sets hybridizing to the middle region (core) and the 5’end of *NEAT1_2* (shell) (**Figure S6A**). We observed two types of PS clusters - the expected elongated cylinder-like structures but also individual PS spheres that were randomly glued together. However, there was no apparent difference in their numbers between SRSF5 KO and WT cells (**Figure S6B**).

Next, to test whether these PS clusters react differently to stress conditions we used four stress conditions known to induce PS clusters ^32^ and first monitored whether the localization of SRSF5 changes. SRSF5-GFP expressing cells were subjected to proteasome inhibition by MG132 (10 µM, 4 h), sodium arsenite (0.25 mM, 1 h), acute hypoxia (24 h 0.2% O2) and the leukemia drug Rocaglamide A (RocA, 5 µM, 4 h; **Figure 7A**). We found that MG132 treatment induced large PS clusters, but they did not co-localize with SRSF5 signal and NS seemed unaffected. In contrast, 1h sodium arsenite treatment completely released SRSF5 from NS, but there was no apparent enrichment of SRSF5 signal in PS. Hypoxia also released SRSF5 from NS, as expected, but here we observed a partial enrichment of SRSF5 around PS. While, RocA treatment, which was shown to clamp RBP binding to purine-rich RNAs ^48^, leaves NS intact, but caused a complete merge of PS clusters and SRSF5 (**Figure 7A**). This merge of PS spheres with SRSF5 speckles was confirmed by super-resolution STED imaging with two different FISH probe sets for *NEAT1_2* (core and shell) (**Figure 7B, S6C-D**). These data show that concentration and release of SRSF5 in NS is stress-regulated and that SRSF5 localizes more to PS clusters in some stress conditions, while it is completely excluded in others suggesting that the shell composition of PS clusters differs between stress conditions.

**Figure 7.**
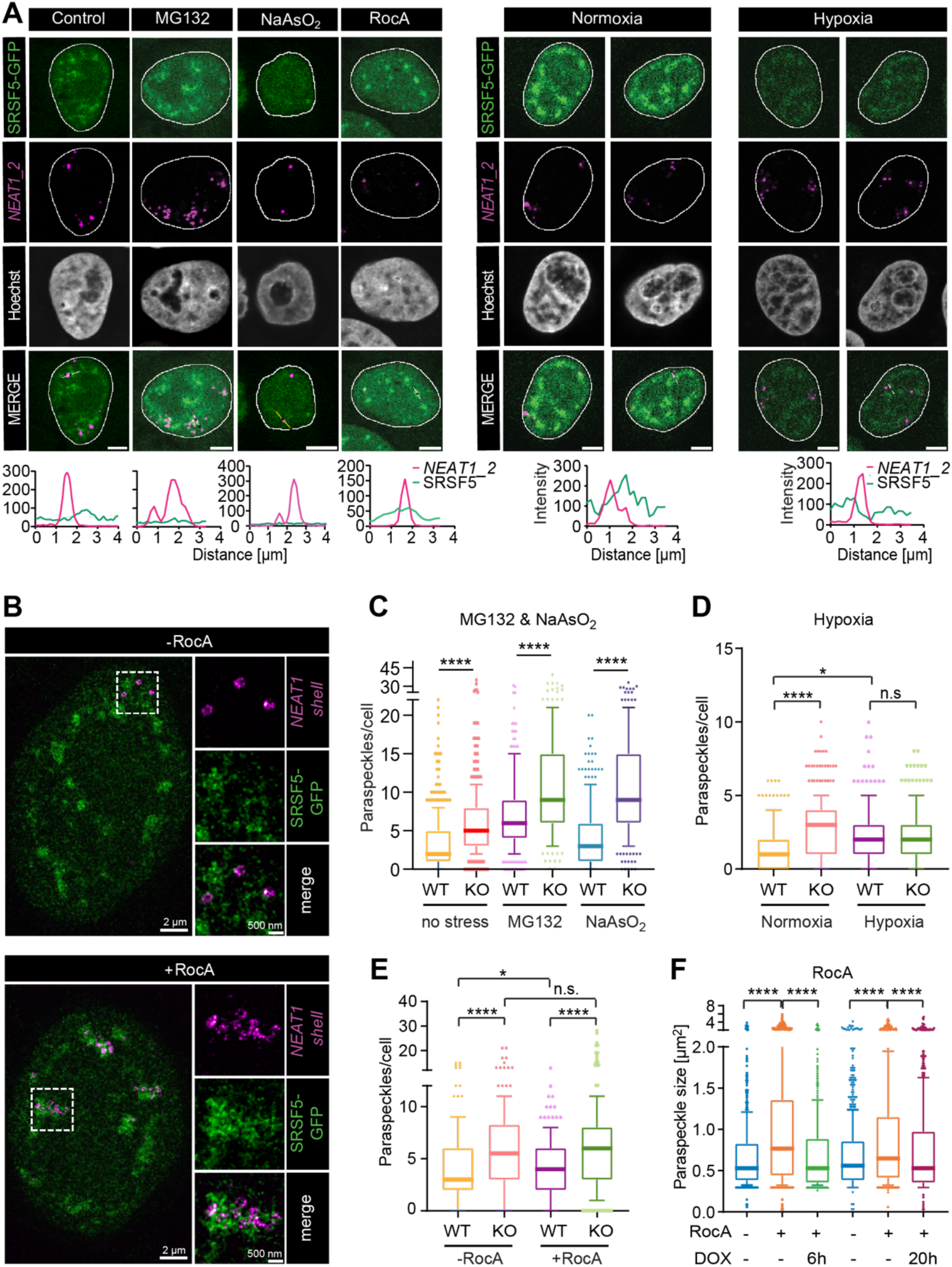
SRSF5 is required for PS hyper-assembly during hypoxia stress and RocA treatment. **A) Top:** Confocal images of HeLa SRSF5-GFP cells subjected to MG132 (4 h, 10 µM), sodium arsenite (4 h, 0.25 mM), RocA (4 h, 5 µM) and hypoxia (24 h, 0.2% O2, 5% CO2). PS were labelled by RNA-FISH targeting the middle region of *NEAT1_2*. Nuclei were stained with Hoechst. Scale bars 5 µm. **Bottom:** Example line scans show partial overlap in control cells, no co-localization in MG132 and sodium arsenite-treated cells and a merge of SRSF5-GFP and *NEAT1_2* signal in hypoxia and RocA treatment. **B)** Quantification of the number of PS spheres per cell from minimum n = 191 to maximum n = 428 cells. Significance was assessed using the Mann-Whitney test. **C)** Superresolution images of Hela SRSF5-GFP cells treated with or without RocA (4 h, 5 µM). DNA-PAINT RNA-FISH using probes at the 5’end of *NEAT1_2* (shell) was coupled with dSTORM imaging for SRSF5-GFP using GFP nanobodies. Scale bars - 2 µm and 500 nm. **D-E)** WT and SRSF5 KO cells were subjected to **D)** hypoxia (24 h, 0.2% O2) and **E)** RocA. Quantification of PS per cell from n = 300-333 cells for hypoxia and n = 308-387 cells for RocA. **F)** SRSF5-GFP cells were treated with RocA and DOX (6 h and 20 h, 1 µg/mL) to deplete SRSF5 simultaneously. Quantification of PS cluster size from n = 439 – 724 PS. All statistics: two-tailed Mann-Whitney Test.

Next we subjected WT and SRSF5 KO cells to these four stress conditions and monitored PS cluster formation. Interestingly, although SRSF5 KO cells showed already much more PS clusters than WT cells, WT and KO responded similarly to MG132 and sodium arsenite treatment, with a strong increase in PS clusters per cell (**Figure 7C, S7A**). This suggests that SRSF5 is not only depleted from these PS clusters, it is also not required for further PS hyper-assembly in these conditions. In contrast, in RocA and hypoxia treatments PS hyper-assembly was completely impaired in SRSF5 KO cells, while WT cells responded to both treatments (**Figure 7D-E, S7B**). To further confirm this dependence on SRSF5 for PS cluster assembly, we used SRSF5 hGRAD cells and treated them simultaneously with DOX (6 h and 20 h) and RocA (4 h). Acute SRSF5 depletion strongly reduced the size of PS clusters induced by RocA treatment compared to control cells indicating that their hyper-assembly required SRSF5 (**Figure 7F, 8A, 8B**).

**Figure 8.**
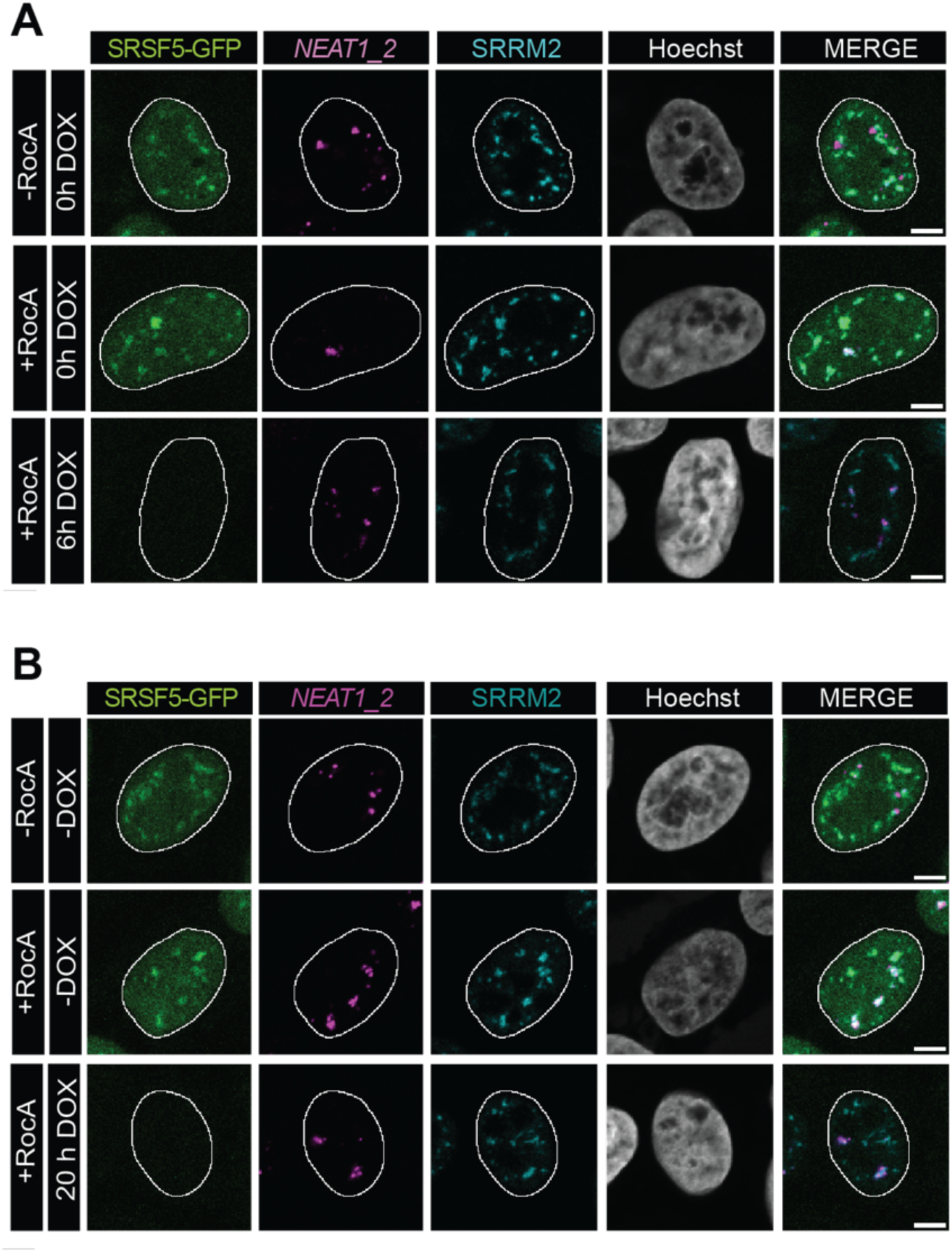
SRSF5 is required for PS hyper-assembly during RocA treatment. Example micrographs of SRSF5-GFP cells treated with RocA (4 h) and DOX (6, 20 h, 1 µg/mL) to deplete SRSF5 simultaneously. PS were labelled by RNA-FISH targeting the middle region of *NEAT1_2*. NS are labelled by IF using an anti-SRRM2 antibody. Nuclei were stained with Hoechst. Scale bars - 5 µm.

Although SRSF5 KO cells did not show any apparent phenotypes nor stress granules (SG) stained by the marker G3BP1 (**Figure S7A-C**), the Nascent-seq data revealed the mis-regulation of many genes involved in intracellular signaling (**Figure S8**), suggesting that even though PS cluster recover in these cells, the stress response is dysregulated in the absence of SRSF5. Indeed, upon stress treatments, SRSF5 KO cells hyper-react and display much more SGs than WT cells (**Figure S7A-C**).

## Discussion

We show here that PS and NS crosstalk and exchange components during cellular homeostasis and stress conditions. Specifically, we found that the nuclear speckle resident protein SRSF5 is dynamically exchanged between both MLOs and is required for PS cluster formation in response to certain stress conditions. Our data suggest that SRSF5 is enriched in PS shells and binds to the 5’end of *NEAT1_2.* Binding promotes the proper alignment of *NEAT1_2* 5’ends to the shell region and the formation of larger PS clusters. The SRSF5 microenvironment might act as glue for individual spheres during PS hyper-assembly (**Figure 3G**). In the absence of SRSF5 *NEAT1_2* is mis-aligned and destabilized, PS are smaller, and cannot form large clusters. While only a small fraction of SRSF5 associates with PS under steady state conditions, this association increases dependent on the type of stress. For example, hypoxia partially disperses NS ^31,49^ and the release of splicing factors allows the reprograming of alternative splicing towards isoforms that promote adaptation to low oxygen. We show here that hypoxia releases SRSF5 from NS and increases its localization to PS. Similarly, after RocA treatment SRSF5 signal in NS is merged with PS. Thus, hypoxia or RocA-specific signaling events regulate NS-PS crosstalk by increasing the possibility for SRSF5 to bind *NEAT1_2* and occupy PS shells, which may contribute to stress adaptation. Interestingly, SRSF5 does not localize to PS in other stress conditions, such as proteasome inhibition and oxidative stress and is also dispensable for PS cluster formation in these conditions. This indicates that there is a stress-specific composition of PS shells likely to ensure an adequate response by sequestering different RNAs and RBPs.

A recent study demonstrated that the 5’end of *NEAT1* and the composition of RBPs that bind to the shell region are important to segregate PS from NS ^28^. It was proposed that RBPs residing in the shell of PS and NS differ fundamentally in their chemical and biophysical properties, e.g. proteins with prion-like domains such as SFPQ versus proteins with mixed charge domains such as SR proteins, and that this increases the energy at the interface between PS and NS ^28^. We identified here the SR protein SRSF5 as a common component of PS and NS that associates with PS shells. But unlike depletion of other shell proteins, including SFPQ, HNRNPF and BRG1, which caused PS and NS to fuse ^28^, acute depletion of SRSF5 did not have this effect. However, treatment with the anti-leukemia drug RocA completely merged SRSF5 and PS. This phenomenon was not described before and the underlying mechanism is unknown. We speculate that this fusion is analog to the artificial tethering of U2snRNP proteins that normally resides in NS, to the PS shell region of *NEAT1* transcript, which caused the internalization of PS within NS ^28^. RocA is known to inhibit translation via clamping of RBPs such as DDX3 and eIF4A to purine-rich RNAs, this way inhibiting their function ^48^. SRSF5 also binds preferentially to purine-rich RNAs and our iCLIP data revealed very high crosslinking of SRSF5 to GA-rich regions at the 5’end of *NEAT1* (not shown). Thus, clamping of SRSF5 and maybe other SR proteins, to purine-rich regions of *NEAT1* could disadvantage the binding of other shell proteins and eventually drag PS into NS. RocA is a known anti-leukemia drug and SRSF5 was recently identified as a pro-leukemia factor ^50^. It would be interesting to study in the future whether RocA’s anti-cancer action is related to SRSF5 inactivation and NS-PS fusion.

It was proposed that the core of PS spheres sequesters components, whereas the shell interacts with different proteins in the nucleoplasm dependent on the function of PS ^4,17^. Our data suggest that different types of PS exist with a distinct shell composition, which might respond differently to stress conditions, given that i) specialized PS containing *NEAT1* were previously reported in the literature ^21,51–53^, ii) only a fraction of SRSF5 localizes to PS, iii) SRSF5 localization changes dramatically dependent on the type of stress, and iv) some of the known PSPs were absent in SRSF5 pulldowns, such as TAF15 and PSPC1, which bind to the shell of PS ^28^ and might compete with SRSF5 for binding sites in *NEAT1*. We assume that in SRSF5 KO cells PS shells are occupied by such competing proteins, which would change the properties of PS spheres. For example, PSPC1 has a prion-like IDR and different biophysical properties and interactors as SRSF5. PSPC1-occupied PS spheres might hyper-assemble in a different way during stress, sequester different RNA and proteins or alter the proximity to NS. As a result, the stress response is de-regulated, as seen in SRSF5 KO cells.

Our data suggest that acute SRSF5 removal destabilizes *NEAT1_2* transcripts. This could be due to its misalignment in PS shells, which may enhance the access of nucleases. However, we had previously suggested a role for SRSF5 in the stabilization of non-coding transcripts through sequestration in NS ^35^. This role of SRSF5 might extend to PS, where SRSF5 creates a protective microenvironment around PS shells.

Using an acute depletion system, we also discovered an intriguing compensation mechanism for PS whereby SRSF5 emerged as a novel regulator of TDP-43 levels. Our data suggest that SRSF5 binding promotes splicing of intron1 and 2 of *TARDBP*, the latter prevents usage of an intronic poly(A)-signal and pre-mature transcription termination. SRSF5 depletion activates intronic polyadenylation and progressively reduces TDP-43 levels. Low TDP43 level promote the switch to the long *NEAT1_2* isoform, which increases PS numbers ^21,52^. In contrast, overexpression of TDP-43, but not a mutant that localizes to the cytoplasm in SRSF5 KO cells reduces PS. This cross-regulatory role of SRSF5 is relevant for example for the understanding of cell fate decisions ^21^. TDP-43 is known to promote pluripotency by repressing the formation of the long *NEAT1_2* isoform. During differentiation TDP-43 levels decrease, *NEAT1_2* is made and PS start to form. These PS sequester existing TDP-43 and inactivate it, which shifts the balance further towards processing of *NEAT1_2* promoting exit from pluripotency. SRSF5 might contribute to this cell fate decision by regulating the levels of TDP-43 during differentiation. We have shown that SRSF5 also promotes the maintenance of pluripotency by exporting transcripts encoding pluripotency factors ^36^. But its export activity is blocked when cells differentiate ^36^. This shuttling switch correlates with the onset of *NEAT1_2* expression. We show here that SRSF5 regulates the levels of TARDBP via splicing, but a decreased export could further reduce the levels of TDP-43 and promote PS formation and exit from pluripotency. Dysregulation of NS or PS dynamics or SRSF5 levels contribute to the development and progression of tumors through the misregulation of alternative splicing and gene expression (REF). Understanding the precise mechanisms for their assembly and crosstalk during stress is crucial for therapeutic intervention and cancer diagnosis. Our work shows that acute depletion systems are essential to distinguish direct from indirect or compensatory effects, as RBPs always act within large compensatory networks. Moreover, we need multi-labelling approaches and very high resolution to quantitatively assess conformational changes or defects in RNA packaging or the architecture of larger PS clusters. Multicolor DNA-PAINT RNA-FISH for super-resolution imaging established in this study, will be a very powerful tool for future studies in this direction.

## Acknowledgements

We are grateful for funding from DAAD to EZ, MMM and DS, CAPES to HYN, the DFG (SFB902-B13 and TRR267-A03 to MMM and KZ, SFB902-A8 to MH) and SCALE (to MMM and MH).

## Author contributions

MMM, BA, LK, EKO and MH designed the experiments. BA, FM, RJR, EKO, LK, HYN and EZ performed and analyzed the experiments. BA, CB, MCHC and KZ performed the bioinformatics analyses. DS and DD contributed plasmids and advice. Figures were prepared by BA, EKO, LK, RJR and MMM. The manuscript was written by BA, EKO, LK, RJR, MH and MMM.

## Material and Methods

### Cell culture, plasmid transfection and drug treatments

HeLa cells were cultivated under humidified condition at 5% CO2 and 37 °C in DMEM GlutaMAX Medium, supplemented with 10% (v/v) heat inactivated fetal bovine serum and 100 µg/mL penicillin-streptomycin (all Gibco™, Thermo Fisher Scientific). For stress experiments, HeLa cells were grown on coverslips placed in 24-well plates and treated with different compounds, followed by fixation with 4% PFA (Thermo Fisher Scientific). The proteasome was inhibited with 10 µM of MG132 (M7449-200UL, Sigma Aldrich), translation was inhibited with 5 µM of RocaglamideA (SML0656-100UG, Sigma Aldrich), and oxidative stress was induced with 0.25 mM of sodium arsenite (041533-AP, Thermo Fisher Scientific). All compounds were diluted in fresh DMEM and cells were treated for 4 h. To induce expression of the hGRAD systems, cells were treated with 1 µg/mL doxycycline (DOX; Sigma-Aldrich, D9891). For hypoxia experiments, coverslips were placed in 6 cm plates and control cells were either grown in normoxic conditions (24 h, 37°C, 0.2% O2, 5% CO2; Hypoxiastation H35, Don Whitely Scientific Limited; HypoxyLab, Oxford Optronics).

Plasmids were extracted from bacteria with the ZR Plasmid Miniprep Classic (Zymo Research). Successful cloning was confirmed by Sanger sequencing (ACGT).

WT HeLa cells were transfected with 0.5 µg purified plasmid DNA per well in 6-well plates using the jetOPTIMUS^®^ (Polyplus-transfection). hGRAD was induced by DOX (1 µg/mL). All plasmids are listed in **Table S7** and all primers in **Table S6**.

### Genome editing with CRISPR/Cas9

SRSF5 KO cells were generated by a dual sgRNA approach. Two sgRNAs were chosen, which cut in the coding sequence within exon2 and exon3 of SRSF5, generating large deletions and frameshifts, subsequently leading to mRNA decay by nonsense-mediated decay. The sgRNAs were annealed from a specific crRNA and a universal tracrRNA (IDT). The crRNAs were designed with the IDT design tool. Site-specific crRNAs were annealed with the universal tracerRNA to sgRNAs and diluted to a final concentration of 1 µM. The crRNA and tracrRNA were first heated to 95°C for 5 minutes and then slowly cooled down to RT to allow optimal sgRNA duplex annealing. Cells were transfected with pre-assembled Cas9-sgRNA RNPs using Lipofectamine CRISPRMAX (Invitrogen) and incubated for another 48 h. Cells were directly used for single-cell generation on 96-well plates. After further incubation (Approximately 7-10 days), all potential single-cell clones were subjected to high throughput genomic screening by PCR and sequencing.

Sequences of the gRNAs are shown in **Table S3** and primers in **Table S6.**

### RNA isolation, reverse transcription and qPCR

Cells were harvested and resuspended in TRIzol™ (Thermo Fisher Scientific). RNA was extracted according to the manufacturer’s instructions, treated with TURBO™ DNase (Thermo Fisher Scientific) for 30 min at 37°C to remove genomic DNA and subsequently purified. Two µg of RNA were reverse transcribed into cDNA using SuperScript™, 10 mM dNTP Mix (both Thermo Fisher Scientific) and oligodT (Sigma Aldrich). qPCR primers were designed using Primer-BLAST (https://www.ncbi.nlm.nih.gov/tools/primer-blast/). qPCRs were performed using cDNA (1:8 dilution) and the ORA™ SEE qPCR Green ROX L kit (highQu) on a PikoReal 96 machine (Thermo Fisher Scientific). GraphPad Prism was used for graphics/statistics. Primers used are listed in **Table S6**.

### Western blot and antibodies

Cells were lysed in 300 µL NET-2 buffer (150 mM NaCl, 0.05% (v/v) NP-40, 50 mM Tris-HCl pH 7.5 supplemented with 1× cOmplete Protease Inhibitor Cocktail (Sigma Aldrich) and 10 mM β-phosphoglycerate (Fluka BioChemica)) or with RIPA buffer (150 mM NaCl, 0.05% (v/v) NP-40, 50 mM Tris-HCl pH 7.5, 0.1% (w/v) SDS, 0.5% (w/v) sodium deoxycholate, freshly added 1× cOmplete Protease Inhibitor Cocktail and 10 mM β-phosphoglycerate). NET-2 lysates were sonicated on ice for 30 sec (3 pulses of 10 sec; 20 sec intervals) at 20% amplitude (Branson W-450 D) and cleared by centrifugation. Protein concentrations were measured using Quick Start Bradford 1× Dye Reagent (Bio-Rad^®^) on a NanoDrop2000 (Thermo Fisher Scientific) or DC Protein-Assay (Bio-Rad^®^) for RIPA samples. 20-40 µg protein were separated by SDS-PAGE on 4-15% Mini-PROTEAN^®^ TGX Stain-Free™ Gels (Bio-Rad^®^) and transferred onto PVDF membranes using Trans-Blot Turbo RTA Mini LF PVDF Transfer Kit (Bio-Rad^®^). Transfer and equal loading were evaluated by activation of stain-free gels by UV light. Membranes were probed with the antibodies listed in **Table S4**. Proteins were imaged using secondary antibodies coupled to a horseradish peroxidase and Amersham™ ECL Prime Western Blotting Detection Reagent (Cytiva) with the ChemiDoc™ MP Imaging System. Image quantification was performed using the ImageLab software (Bio-Rad).

### Nascent-RNA sequencing

HeLa WT and SRSF5 KO cells were incubated with 400 µM 4-thio-uridine (4sU) for 1 h to label newly transcribed RNAs. The labelling reaction was stopped by adding TRIzol™ reagent to cells in the culture dishes. RNA extraction and pull-down of newly transcribed RNA was performed according to (Gressel et al., 2019) without fractionation of the RNA by sonification. Nascent RNA was purified using RNA Clean and Concentrate Kit (ZymoResearch). cDNA libraries for RNA-seq were prepared with universal Plus™ Total RNA-Seq Library Preparation Kit (Tecan) according to manufacturer’s instructions. Ribosomal RNA fragments were removed and the library was sequenced on an Illumina NovaSeq6000 instrument (2 replicates per time point, 100 Mio reads, 150 bp, paired-end). Reads were mapped to the human genome (hg38) using the STAR mapper (version 2.7.10) (Dobin et al., 2013) with the following parameters: --outFilterMultimapNmax 1 -- outFilterMismatchNmax 999 --outFilterMismatchNoverReadLmax 0.04 --outSAMtype BAM SortedByCoordinate.

Differential gene expression was quantified using DESeq2 with default parameters (Love et al., 2014). Gene Ontology (GO) enrichment analysis was performed with hypeR or with the over-representation test implemented in the enrichGO function of the clusterProfiler package in R (Yu et al., 2012). Enrichment was tested against the union of all genes that were tested in the DESeq2 analysis. Adjusted *P* value cutoffs were set to 0.05 and “Biological Process” categories were explored.

Splicing analysis was performed using MAJIQ (version 2.2) (Vaquero-Garcia et al., 2016). Specifically, we used the majiq-build function to construct a single splice graph over all replicates and conditions (WT, SRSF5 KO, T0, T2, T8 and T16). Next, majiq-deltapsi was used to quantify local splicing variations (LSVs) between WT and SRSF5 KO and T0, T2, T8 and T16 datasets. Identified LSVs from both comparisons were considered significantly regulated with a difference in junction usage (percent selected index, PSI) of DPSI > 5% and a changing probability > 50%. Non-regulated LSVs were considered with DPSI < 2% and an LSV changing probability of 0%. Both regulated and non-regulated LSVs were subsequently stratified to the binary level. First, we identified the two main junctions per LSV based on the DPSI, retaining only those LSVs were at least 50% of the change seen in the strongest junction can be explained by the second strongest junction. LSVs were classified into the events ‘intron-retention’, ‘alternative 3’ splice site’, ‘alternative 5’ splice site’ and ‘cassette exon’. This was done by combining source and target LSV junctions for a given event, requiring both LSVs to be regulated as defined above. Cassette exon events were further sub-classified into ‘simple cassette exon’, ‘alternative first exon’, ‘alternative last exon’ and ‘complex events’ using the same methodology.

### Confocal image acquisition and quantification

Images were acquired with confocal laser-scanning microscope (LSM780; ZEISS) using the Zen 2012 (black edition; 8.0.5.273; ZEISS). Fluorescence signal was detected with an Argon laser, (GFP – 488nm, Qasar 570 – 561nm and Qasar 670 – 647nm). Images from the same experiment were acquired with the same settings for all conditions. Line scans were performed by drawing a straight line across the nucleus, without crossing the nucleolus. Fluorescence intensity per pixel in the line area was acquired using the “Plot Profile” tool. Images were analyzed using Fiji (Schindelin et al., 2012). Pictures were cropped with the Image crop function and scale bars were added. For fluorescence quantification, the Hoechst channel was used to acquire a threshold image (“Threshold Li”) and the “Particle analyzer” plug-in from the Biovoxxel toolbox (https://imagej.net/BioVoxxel_Toolbox) was used to obtain the nuclear regions of interest (ROI). ROIs were transferred to the GFP channel and fluorescence was quantified using the ‘integrated density value’ (mean gray value per pixel x area). Data were plotted using GraphPad Prism 8 (https://www.graphpad.com), and cell lines and conditions were compared using the Mann-Whitney test.

### Nanobody labeling

Unlabelled nanobody against GFP (anti-GFP single domain Antibody, #N0305-250µg) with a single ectopic cysteine at the C terminus was ordered from NanoTag Biotechnologies (Germany). Nanobodies were self-labelled with azide-modified docking strands according to a previously published protocol ^54^. In brief, 20 nmol of nanobodies in 1x phosphate buffer saline (PBS) were incubated with a final concentration of 5 mM TCEP (Sigma-Aldrich, Netherlands) for 2 h at 4 °C to reduce the ectopic cysteine. Afterwards, the excess of TCEP was removed by exchanging the buffer to PBS pH 6.5 using Amicon spin filter (MWCO 10 kDa, Merck, Germany). The DBCO-sulfo-NHS ester linker (Jena Bioscience, Germany) was dissolved in dimethylformamide (DMF) (Sigma-Aldrich, Germany) and then diluted in 1x PBS. Linker and nanodbody were mixed in a 10:1 molar ratio and incubated for 90 min at 4°C gently shaking. The unbound linker was removed using Amicon spin filter (MWCO 10 kDa). Azide-modified P1 docking strand (**Table S2**) and linker-nanobody conjugate were incubated in a 10:1 molar ratio overnight at 4°C while slightly shaking. The next day, unbound DNA was removed using an Amicon spin filter (MWCO 10 kDa). The docking strand-labelled nanobody was stored at 4°C.

### RNA Fluorescence in Situ Hybridization (FISH) and Immunofluorescence (IF)

For FISH-IF experiments, 12-mm coverslips were placed inside 10-cm plates used for the experiments. After removing the medium and washing the cells with 1× PBS, the coverslips were transferred to 24-well plates. Cells were fixed with 4% PFA in PBS for 15 min, washed with PBS and permeabilized with 70% ethanol for 1 h. FISH was performed using Stellaris probes and buffers (LG Biosearch Technologies) following the manufacturer’s protocol. Coverslips were washed with Stellaris Wash Buffer A followed by incubation in Stellaris Hybridization buffer containing the FISH probes and the primary antibodies, placed in a humidified chamber and hybridized for 16 h to 20 h at 37°C protected from light. A mouse α -SRRM2 antibody (PA5-59559, 1:70 dilution in Stellaris Hybridization buffer) was used to detect nuclear speckles. To visualize stress granules, mouse α-G3BP1 antibody (ab56574, 2 µg/mL final concentration in Stellaris Hybridization buffer) was used as a marker. After hybridization, the coverslips were incubated with Stellaris Wash Buffer A containing the secondary antibody (Goat anti-Rabbit IgG Alexa Fluor™ Plus 555, ThermoFisher **A32732**; Goat anti-Rabbit IgG Alexa Fluor™ 488, ThermoFisher A-11008; Donkey anti-Mouse IgG Alexa Fluor™ 594, ThermoFisher **A-21203;** Donkey anti-Rabbit IgG Alexa Fluor™ 594, ThermoFisher A-21207) in a 1:500 dilution for 30 min at 37°C. DNA was stained with Hoechst 34580 (Sigma Aldrich) at a final concentration of 0,25 µg/mL in Wash Buffer A for 30 min at 37°C. The coverslips were washed with Wash Buffer B, dried for 15 min and mounted onto glass slides using ProLong Diamond Antifade Mountant (Invitrogen). FISH probes used are listed in **Table S1** and antibodies in **Table S4**.

For super-resolved immunofluorescence fixed cells were blocked using immunofluorescence staining (IF) buffer (3% (w/v) BSA, 0.1% (v/v) Triton-X 100 in 1x PBS) for 30 min and in the following incubated with the self-labelled anti-GFP nanobodies at a concentration of approximately 50 nM in IF buffer for 1.5 h at room temperature while shaking. Afterwards the samples were washed three times with 1x PBS and then fixed with 4% (v/v) formaldehyde for 10 min at room temperature and washed three times with 1x PBS. Gold beads (Gold nanoparticles, 100 nm diameter, Product #A11-100-NPC-DIH-1-25, NanoPartz, USA) as fiducial markers were diluted 1:5 in 1x PBS and incubated for 10 min.

### dSTORM imaging

*d*STORM imaging of HeLa cells was carried out on a home-built widefield setup based on a Nikon Eclipse Ti microscope ^37^. The excitation light was generated by a DPSS laser at 640 nm (LPX-640L-500-CSB-PPA, Oxxius S.A, France) and a laser diode at 488 nm (LBX-488-200-CSB-PPA, Oxxius S.A, France) with the required excitation power controlled by an acousto-optic tunable filter (AOTFnC-400.650-TN, AA Opto Electronic, France). To clean the beam-profile the laser was coupled by a fibre collimator (60FC-4-M6.2-33, Schäfter & Kirchhoff GmbH, Germany) into a polarisation maintaining single-mode optical fibre (PMC-E-400RGB, Schäfter & Kirchhoff GmbH, Germany) and subsequently recollimated to a FWHM diameter of 6 mm (60FC-T-4-M50L-01, Schäfter & Kirchhoff GmbH, Germany). The collinear beam was then directed through two telescope lenses (AC255-030-A-ML and AC508-150-A-ML, Thorlabs GmbH, Germany) which focused the beam onto the back focal plane of the objective (CFI Apochromat TIRF 100XC Oil, NA 1.49, Nikon, Japan). A mirror mounted on a motorised translation stage (MTS50-Z8, Thorlabs GmbH, Germany) was used to vary the illumination angle between widefield, HILO or TIRF. The excitation light was coupled into the microscope by means of a dielectric beamsplitter (zt405/488/561/640rpc, AHF Analysentechnik, Germany) which also transmitted the emission light into the detection beam path. The axial focus was maintained using an autofocus system (Ti-PFS, Nikon) and the lateral position was adjusted using a motorised stage (Ti-S-ER, Nikon) combined with a piezo stage (Nano-Drive, MadCityLabs, USA). After spectral filtering with a bandpass filter (700/75 ET, Chroma Technology Corp., USA) the emission light was projected onto an Andor Ixon Ultra EMCCD camera (DU-897U-CS0, Andor, North Ireland). *NEAT1* samples were excited with a laser intensity of 135 W/cm² and SRSF5 with 215 W/cm² in HILO illumination. *d*STORM imaging of *NEAT1* was performed in *d*STORM imaging buffer (1x PBS, 100 mM β-mercaptoethylamine, 50 U/mL glucose oxidase, 5000 U/mL catalase, 0.2 mM Tris (2-carboxylethyl) phosphine hydrochloride, 20% (w/v) glucose, pH = 8.5). Before the start of the measurement, the sample was already exposed for 7.5 min to allow the equilibrium to be set. A preamplifier gain of 3 and an EM-gain of 200 were used to record 35,000 frames with an integration time of 30 ms.

### dSTORM data post processing

*d*STORM data was post processed with the Picasso software (v0.6.0) ^38^. Localization of single molecule signals was performed using Picasso Localise module. The following parameters were used: baseline 82.3 photons, sensitivity 5.14, quantum efficiency 0.95, pixel size 158 nm and a min net gradient of 8,000 for *NEAT1* and 10,000 for SRSF5. Reconstruction of the super-resolution images was performed using Picasso Render module. The images were drift corrected with redundant cross correlation (RCC) and stacks of 2000-4000 frames. Two color images were aligned using gold beads as fiducial markers (Gold nanoparticles, 100 nm diameter, Product #A11-100-NPC-DIH-1-25, NanoPartz, USA). Localisations from *NEAT1* were filtered for the ellipticity (0-0.15) and the width of the point spread function (0.8-1.15 px) and linked with 4x NeNA and 2 transient dark frames. Localisations from SRSF5 were filtered for the ellipticity (0 0.2) and the width of the point spread function (0.7-1.3 px) and linked with 4x NeNA and 6 transient dark frames. Images were exported with a 10 nm pixel size.

### dSTORM data analysis

For the threshold-based segmentation of the paraspeckle signal, custom-written Fiji macros were used. Therefore, the super-resolved images were duplicated, smoothed with a Gaussian blur of 10 nm and then converted into a binary mask. The necessary threshold was first tested manually and then applied to all measurements. The objects were then counted and measured automatically with the “Particle analyzer” plug-in from the Biovoxxel toolbox (https://imagej.net/BioVoxxel_Toolbox). To quantify the frequency and size of individual paraspeckles, the segmented objects were filtered by the following parameters: size (>0.025 µm²) and roundness (>0.8). Roundness is defined as:

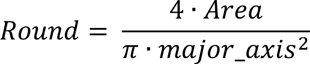

Size (>0.2 µm²) was filtered to quantify paraspeckle clusters. Two-sample t- and Mann-Whitney-U-tests were performed using GraphPad PRISM (GraphPad Software, USA).

### STED imaging

Two-color STED imaging of *NEAT1* was performed using an Abberior Expert Line microscope (Abberior Instruments, Germany) equipped with an Olympus IX83 stand (Olympus Deutschland GmbH, Germany) and a UPLXAPO 60x NA 1.42 oil immersion objective (Olympus Deutschland GmbH, Germany). For image acquisition, the sample was excited with a 561-nm and a 640-nm excitation laser (1.5 µW and 16 µW respectively at the back focal plane) and depleted with a pulsed 775-nm laser (506 mW and 17 mW at the back focal plane) using a 2D donut point broadening function and a delay of 750 ps, and a detection width of 8 ns. Fluorescence signals were acquired in the spectral regions from 571 nm to 630 nm and from 650 nm to 760 nm using two avalanche photodiode (APD) detectors. Images were acquired with a pinhole aperture of 0.81 AU, a line accumulation of 60, a pixel dwell time of 5 µs, and a pixel size of 20×20 nm². Measurements were acquired in line sequential mode. For STED-PAINT of *NEAT1*, imager strands (**Table S1**) were diluted to a concentration of 100 nM in DNA-PAINT imaging buffer (1x PBS, 0.5 M NaCl).

### STED data quantification

The analysis of the two-colored paraspeckles in the STED images was performed with the help of a self-written MATLAB (MathWorks, USA) script. In summary, the round signals of the 5’ probes were segmented using the Hough transformation with a sensitivity of 0.85 and then matched to the corresponding 3’ signals. The segmented signals were then fitted using a ring function:

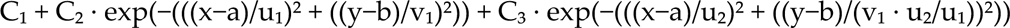

The distance between the center of the ring and the outer FWHM was taken as the radius of the ring-shaped 5’ or 3’ signal.

## References

1. Adriaens, C., and Marine, J.C. (2017). NEAT1-containing paraspeckles: Central hubs in stress response and tumor formation. Cell Cycle 16, 137–138. 10.1080/15384101.2016.1235847.

2. Hirose, T., Ninomiya, K., Nakagawa, S., and Yamazaki, T. (2023). A guide to membraneless organelles and their various roles in gene regulation. Nat Rev Mol Cell Biol 24, 288–304. 10.1038/s41580-022-00558-8.

3. Staněk, D., and Fox, A.H. (2017). Nuclear bodies: news insights into structure and function. Curr Opin Cell Biol 46, 94–101. 10.1016/j.ceb.2017.05.001.

4. McCluggage, F., and Fox, A.H. (2021). Paraspeckle nuclear condensates: Global sensors of cell stress? Bioessays 43, e2000245. 10.1002/bies.202000245.

5. Clemson, C.M., Hutchinson, J.N., Sara, S.A., Ensminger, A.W., Fox, A.H., Chess, A., and Lawrence, J.B. (2009). An architectural role for a nuclear noncoding RNA: NEAT1 RNA is essential for the structure of paraspeckles. Mol Cell 33, 717–726. 10.1016/j.molcel.2009.01.026.

6. Hutchinson, J.N., Ensminger, A.W., Clemson, C.M., Lynch, C.R., Lawrence, J.B., and Chess, A. (2007). A screen for nuclear transcripts identifies two linked noncoding RNAs associated with SC35 splicing domains. BMC Genomics 8, 39. 10.1186/1471-2164-8-39.

7. Sasaki, Y.T., Ideue, T., Sano, M., Mituyama, T., and Hirose, T. (2009). MENepsilon/beta noncoding RNAs are essential for structural integrity of nuclear paraspeckles. Proc Natl Acad Sci U S A 106, 2525–2530. 10.1073/pnas.0807899106.

8. Sunwoo, H., Dinger, M.E., Wilusz, J.E., Amaral, P.P., Mattick, J.S., and Spector, D.L. (2009). MEN epsilon/beta nuclear-retained non-coding RNAs are up-regulated upon muscle differentiation and are essential components of paraspeckles. Genome Res 19, 347–359. 10.1101/gr.087775.108.

9. Hirose, T., Yamazaki, T., and Nakagawa, S. (2019). Molecular anatomy of the architectural NEAT1 noncoding RNA: The domains, interactors, and biogenesis pathway required to build phase-separated nuclear paraspeckles. Wiley Interdiscip Rev RNA 10, e1545. 10.1002/wrna.1545.

10. Kawaguchi, T., Tanigawa, A., Naganuma, T., Ohkawa, Y., Souquere, S., Pierron, G., and Hirose, T. (2015). SWI/SNF chromatin-remodeling complexes function in noncoding RNA-dependent assembly of nuclear bodies. Proc Natl Acad Sci U S A 112, 4304–4309. 10.1073/pnas.1423819112.

11. Naganuma, T., Nakagawa, S., Tanigawa, A., Sasaki, Y.F., Goshima, N., and Hirose, T. (2012). Alternative 3’-end processing of long noncoding RNA initiates construction of nuclear paraspeckles. EMBO J 31, 4020–4034. 10.1038/emboj.2012.251.

12. Yamazaki, T., Souquere, S., Chujo, T., Kobelke, S., Chong, Y.S., Fox, A.H., Bond, C.S., Nakagawa, S., Pierron, G., and Hirose, T. (2018). Functional Domains of NEAT1 Architectural lncRNA Induce Paraspeckle Assembly through Phase Separation. Mol Cell 70, 1038–1053.e1037. 10.1016/j.molcel.2018.05.019.

13. Hennig, S., Kong, G., Mannen, T., Sadowska, A., Kobelke, S., Blythe, A., Knott, G.J., Iyer, K.S., Ho, D., Newcombe, E.A., et al. (2015). Prion-like domains in RNA binding proteins are essential for building subnuclear paraspeckles. J Cell Biol 210, 529–539. 10.1083/jcb.201504117.

14. Wang, Y., and Chen, L.L. (2020). Organization and function of paraspeckles. Essays Biochem 64, 875–882. 10.1042/EBC20200010.

15. Chujo, T., and Hirose, T. (2017). Nuclear Bodies Built on Architectural Long Noncoding RNAs: Unifying Principles of Their Construction and Function. Mol Cells 40, 889–896. 10.14348/molcells.2017.0263.

16. Souquere, S., Beauclair, G., Harper, F., Fox, A., and Pierron, G. (2010). Highly ordered spatial organization of the structural long noncoding NEAT1 RNAs within paraspeckle nuclear bodies. Mol Biol Cell 21, 4020–4027. 10.1091/mbc.E10-08-0690.

17. West, J.A., Mito, M., Kurosaka, S., Takumi, T., Tanegashima, C., Chujo, T., Yanaka, K., Kingston, R.E., Hirose, T., Bond, C., et al. (2016). Structural, super-resolution microscopy analysis of paraspeckle nuclear body organization. J Cell Biol 214, 817–830. 10.1083/jcb.201601071.

18. Yamazaki, T., Fujikawa, C., Kubota, A., Takahashi, A., and Hirose, T. (2018). CRISPRa-mediated NEAT1 lncRNA upregulation induces formation of intact paraspeckles. Biochem Biophys Res Commun 504, 218–224. 10.1016/j.bbrc.2018.08.158.

19. Wilusz, J.E., JnBaptiste, C.K., Lu, L.Y., Kuhn, C.D., Joshua-Tor, L., and Sharp, P.A. (2012). A triple helix stabilizes the 3’ ends of long noncoding RNAs that lack poly(A) tails. Genes Dev 26, 2392–2407. 10.1101/gad.204438.112.

20. Barra, J., Gaidosh, G.S., Blumenthal, E., Beckedorff, F., Tayari, M.M., Kirstein, N., Karakach, T.K., Jensen, T.H., Impens, F., Gevaert, K., et al. (2020). Integrator restrains paraspeckles assembly by promoting isoform switching of the lncRNA. Sci Adv 6, eaaz9072. 10.1126/sciadv.aaz9072.

21. Modic, M., Grosch, M., Rot, G., Schirge, S., Lepko, T., Yamazaki, T., Lee, F.C.Y., Rusha, E., Shaposhnikov, D., Palo, M., et al. (2019). Cross-Regulation between TDP-43 and Paraspeckles Promotes Pluripotency-Differentiation Transition. Mol Cell 74, 951–965.e913. 10.1016/j.molcel.2019.03.041.

22. Yamazaki, T., Yamamoto, T., Yoshino, H., Souquere, S., Nakagawa, S., Pierron, G., and Hirose, T. (2021). Paraspeckles are constructed as block copolymer micelles. EMBO J 40, e107270. 10.15252/embj.2020107270.

23. Yamazaki, T., Yamamoto, T., and Hirose, T. (2022). Micellization: A new principle in the formation of biomolecular condensates. Front Mol Biosci 9, 974772. 10.3389/fmolb.2022.974772.

24. Ilık, İ., and Aktaş, T. (2022). Nuclear speckles: dynamic hubs of gene expression regulation. FEBS J 289, 7234–7245. 10.1111/febs.16117.

25. Fei, J., Jadaliha, M., Harmon, T.S., Li, I.T.S., Hua, B., Hao, Q., Holehouse, A.S., Reyer, M., Sun, Q., Freier, S.M., et al. (2017). Quantitative analysis of multilayer organization of proteins and RNA in nuclear speckles at super resolution. J Cell Sci 130, 4180–4192. 10.1242/jcs.206854.

26. Liao, S.E., and Regev, O. (2021). Splicing at the phase-separated nuclear speckle interface: a model. Nucleic Acids Res 49, 636–645. 10.1093/nar/gkaa1209.

27. Paul, S., Arias, M.A., Wen, L., Liao, S.E., Zhang, J., Wang, X., Regev, O., and Fei, J. (2024). RNA molecules display distinctive organization at nuclear speckles. iScience 27, 109603. 10.1016/j.isci.2024.109603.

28. Takakuwa, H., Yamazaki, T., Souquere, S., Adachi, S., Yoshino, H., Fujiwara, N., Yamamoto, T., Natsume, T., Nakagawa, S., Pierron, G., and Hirose, T. (2023). Shell protein composition specified by the lncRNA NEAT1 domains dictates the formation of paraspeckles as distinct membraneless organelles. Nat Cell Biol 25, 1664–1675. 10.1038/s41556-023-01254-1.

29. Faraway, R., Heaven, N.C., Digby, H., Wilkins, O.G., Chakrabarti, A.M., Iosub, I.A., Knez, L., Ameres, S.L., Plaschka, C., and Ule, J. (2023). Mutual homeostasis of charged proteins. bioRxiv, 2023.2008.2021.554177. 10.1101/2023.08.21.554177.

30. Müller-McNicoll, M., Botti, V., de Jesus Domingues, A.M., Brandl, H., Schwich, O.D., Steiner, M.C., Curk, T., Poser, I., Zarnack, K., and Neugebauer, K.M. (2016). SR proteins are NXF1 adaptors that link alternative RNA processing to mRNA export. Genes Dev 30, 553–566. 10.1101/gad.276477.115.

31. de Oliveira Freitas Machado, C., Schafranek, M., Brüggemann, M., Hernández Cañás, M.C., Keller, M., Di Liddo, A., Brezski, A., Blümel, N., Arnold, B., Bremm, A., et al. (2023). Poison cassette exon splicing of SRSF6 regulates nuclear speckle dispersal and the response to hypoxia. Nucleic Acids Res 51, 870–890. 10.1093/nar/gkac1225.

32. An, H., Tan, J.T., and Shelkovnikova, T.A. (2019). Stress granules regulate stress-induced paraspeckle assembly. J Cell Biol 218, 4127–4140. 10.1083/jcb.201904098.

33. Cooper, D.R., Carter, G., Li, P., Patel, R., Watson, J.E., and Patel, N.A. (2014). Long Non-Coding RNA NEAT1 Associates with SRp40 to Temporally Regulate PPARγ2 Splicing during Adipogenesis in 3T3-L1 Cells. Genes (Basel) 5, 1050–1063. 10.3390/genes5041050.

34. West, J.A., Davis, C.P., Sunwoo, H., Simon, M.D., Sadreyev, R.I., Wang, P.I., Tolstorukov, M.Y., and Kingston, R.E. (2014). The long noncoding RNAs NEAT1 and MALAT1 bind active chromatin sites. Mol Cell 55, 791–802. 10.1016/j.molcel.2014.07.012.

35. Arnold, B., Riegger, R.J., Okuda, E.K., Slišković, I., Keller, M., Bakisoglu, C., McNicoll, F., Zarnack, K., and Müller-McNicoll, M. (2024). hGRAD: A versatile “one-fits-all” system to acutely deplete RNA binding proteins from condensates. J Cell Biol 223. 10.1083/jcb.202304030.

36. Botti, V., McNicoll, F., Steiner, M.C., Richter, F.M., Solovyeva, A., Wegener, M., Schwich, O.D., Poser, I., Zarnack, K., Wittig, I., et al. (2017). Cellular differentiation state modulates the mRNA export activity of SR proteins. J Cell Biol 216, 1993–2009. 10.1083/jcb.201610051.

37. Kessler, L.F., Balakrishnan, A., Deußner-Helfmann, N.S., Li, Y., Mantel, M., Glogger, M., Barth, H.D., Dietz, M.S., and Heilemann, M. (2023). Self-quenched Fluorophore Dimers for DNA-PAINT and STED Microscopy. Angew Chem Int Ed Engl 62, e202307538. 10.1002/anie.202307538.

38. Schnitzbauer, J., Strauss, M.T., Schlichthaerle, T., Schueder, F., and Jungmann, R. (2017). Super-resolution microscopy with DNA-PAINT. Nat Protoc 12, 1198–1228. 10.1038/nprot.2017.024.

39. Hell, S.W., and Wichmann, J. (1994). Breaking the diffraction resolution limit by stimulated emission: stimulated-emission-depletion fluorescence microscopy. Opt Lett 19, 780–782. 10.1364/ol.19.000780.

40. Glogger, M., Spahn, C., Enderlein, J., and Heilemann, M. (2021). Multi-Color, Bleaching-Resistant Super-Resolution Optical Fluctuation Imaging with Oligonucleotide-Based Exchangeable Fluorophores. Angew Chem Int Ed Engl 60, 6310–6313. 10.1002/anie.202013166.

41. Spahn, C., Hurter, F., Glaesmann, M., Karathanasis, C., Lampe, M., and Heilemann, M. (2019). Protein-Specific, Multicolor and 3D STED Imaging in Cells with DNA-Labeled Antibodies. Angew Chem Int Ed Engl 58, 18835–18838. 10.1002/anie.201910115.

42. Schwalb, B., Michel, M., Zacher, B., Frühauf, K., Demel, C., Tresch, A., Gagneur, J., and Cramer, P. (2016). TT-seq maps the human transient transcriptome. Science 352, 1225–1228. 10.1126/science.aad9841.

43. Love, M.I., Huber, W., and Anders, S. (2014). Moderated estimation of fold change and dispersion for RNA-seq data with DESeq2. Genome Biol 15, 550. 10.1186/s13059-014-0550-8.

44. Isobe, M., Toya, H., Mito, M., Chiba, T., Asahara, H., Hirose, T., and Nakagawa, S. (2020). Forced isoform switching of Neat1_1 to Neat1_2 leads to the loss of Neat1_1 and the hyperformation of paraspeckles but does not affect the development and growth of mice. RNA 26, 251–264. 10.1261/rna.072587.119.

45. Kawachi, T., Masuda, A., Yamashita, Y., Takeda, J.I., Ohkawara, B., Ito, M., and Ohno, K. (2021). Regulated splicing of large exons is linked to phase-separation of vertebrate transcription factors. EMBO J 40, e107485. 10.15252/embj.2020107485.

46. Schwich, O.D., Blümel, N., Keller, M., Wegener, M., Setty, S.T., Brunstein, M.E., Poser, I., Mozos, I.R.L., Suess, B., Münch, C., et al. (2021). SRSF3 and SRSF7 modulate 3’UTR length through suppression or activation of proximal polyadenylation sites and regulation of CFIm levels. Genome Biol 22, 82. 10.1186/s13059-021-02298-y.

47. Jha, A., Gazzara, M.R., and Barash, Y. (2017). Integrative deep models for alternative splicing. Bioinformatics 33, i274–i282. 10.1093/bioinformatics/btx268.

48. Chen, M., Asanuma, M., Takahashi, M., Shichino, Y., Mito, M., Fujiwara, K., Saito, H., Floor, S.N., Ingolia, N.T., Sodeoka, M., et al. (2021). Dual targeting of DDX3 and eIF4A by the translation inhibitor rocaglamide A. Cell Chem Biol 28, 475–486.e478. 10.1016/j.chembiol.2020.11.008.

49. McIntyre, A.B.R., Tschan, A.B., Meyer, K., Walser, S., Rai, A.K., Fujita, K., and Pelkmans, L. (2024). Phosphorylation-controlled cohesion of a nuclear condensate regulates mRNA retention. bioRxiv, 2023.2008.2021.554101. 10.1101/2023.08.21.554101.

50. Chang, J., Yan, S., Geng, Z., and Wang, Z. (2023). Inhibition of splicing factors SF3A3 and SRSF5 contributes to As. Arch Biochem Biophys 743, 109677. 10.1016/j.abb.2023.109677.

51. Wang, C., Duan, Y., Duan, G., Wang, Q., Zhang, K., Deng, X., Qian, B., Gu, J., Ma, Z., Zhang, S., et al. (2020). Stress Induces Dynamic, Cytotoxicity-Antagonizing TDP-43 Nuclear Bodies via Paraspeckle LncRNA NEAT1-Mediated Liquid-Liquid Phase Separation. Mol Cell 79, 443–458.e447. 10.1016/j.molcel.2020.06.019.

52. Shelkovnikova, T.A., Kukharsky, M.S., An, H., Dimasi, P., Alexeeva, S., Shabir, O., Heath, P.R., and Buchman, V.L. (2018). Protective paraspeckle hyper-assembly downstream of TDP-43 loss of function in amyotrophic lateral sclerosis. Mol Neurodegener 13, 30. 10.1186/s13024-018-0263-7.

53. Zhao, J., Xie, W., Yang, Z., Zhao, M., Ke, T., Xu, C., Li, H., Chen, Q., and Wang, Q.K. (2022). Identification and characterization of a special type of subnuclear structure: AGGF1-coated paraspeckles. FASEB J 36, e22366. 10.1096/fj.202101690RR.

54. Sograte-Idrissi, S., Oleksiievets, N., Isbaner, S., Eggert-Martinez, M., Enderlein, J., Tsukanov, R., and Opazo, F. (2019). Nanobody Detection of Standard Fluorescent Proteins Enables Multi-Target DNA-PAINT with High Resolution and Minimal Displacement Errors. Cells 8. 10.3390/cells8010048.

