## Supplemental material for "Rapid depletion and super-resolution microscopy reveal an unexpected role of the nuclear-speckle protein SRSF5 in paraspeckle assembly and dynamics during cellular stress"

**Formatted:** Font colour: Text 1

**Deleted:** Benjamin Arnold[1,§], Laurell Kessler[2,§], Ellen Kazumi Okuda[1,3,§], Ricarda J. Riegger[1], Maria Clara Hernández Cañás[1,4], Ewelina Zebrowska[1], Cem Bakisoglu[1,4], David Stanek[5], Dorothee Dormann[6], Kathi Zarnack[1,4], Mike Heilemann[2,#] & Michaela Müller-McNicoll[1,7#]

[1]Institute of Molecular Biosciences, Goethe University, Frankfurt am Main, Germany

[2]Institute of Physical and Theoretical Chemistry, Goethe University, Frankfurt am Main, Germany

[3]IMPRS on Cellular Biophysics

[4]Buchmann Institute for Molecular Life Sciences (BMLS), Frankfurt am Main, Germany

[5]Institute of Molecular Genetics (IMG), CAS, Prague, Czech Republic

[6]Institute for Molecular Physiology, Gutenberg University, Mainz, Germany

[7]Max Planck Institute for Biophysics, Frankfurt am Main, Germany

### SUPPLEMENTARY MATERIAL

Content:

Supplementary Figures ........................................................................................................2

Supplementary Tables........................................................................................................13

#### Supplementary Figures

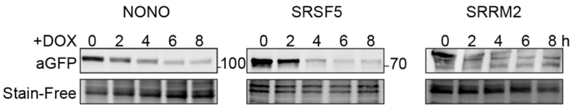


**Figure S1: Acute depletion of SRSF5 decreases paraspeckle number and size. A)** Representative Western blots of the degradation timeline of NONO-, SRSF5- and SRRM2-GFP after induction of hGRAD by DOX (1 µg/mL) for 8 h. Stain-free membranes were used to control for equal loading

.

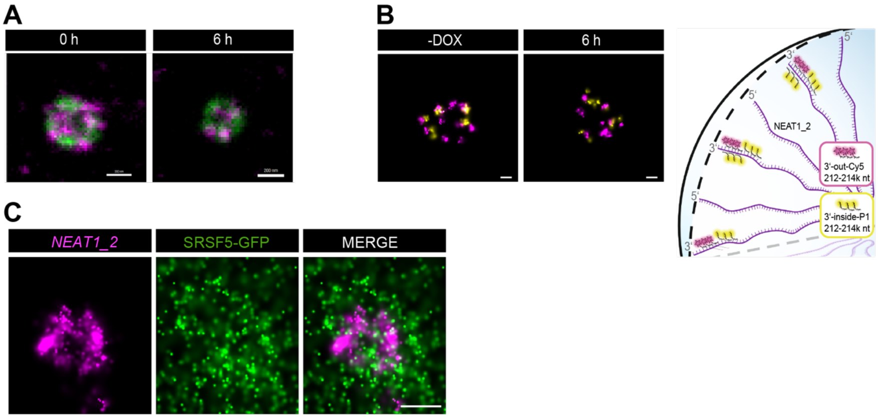


**Figure S2. Paraspeckles are smaller in diameter and differently packaged in the absence of SRSF5. A)** Multicolor DNA-PAINT RNA-FISH was used to label 5' and 3'end of *NEAT1_2*. dSTORM image of one representative PS sphere at 0 h at 6 h SRSF5 depletion. Scale bars - 200 nm. **B)** Multicolor DNA-PAINT RNA-FISH was used to label two neighboring regions at the 3'end of *NEAT1_2*. **Left:** dSTORM image of one representative PS sphere at 0 h at 6 h SRSF5 depletion. Scale bars - 100 nm. **Right:** Positions of where the probes bind to the *NEAT1_2* RNA are indicated. **C)** DNA-PAINT RNA-FISH using probes at the 5'end of *NEAT1_2* was coupled with dSTORM imaging for SRSF5-GFP using GFP nanobodies. Scale bars - 200 nm.

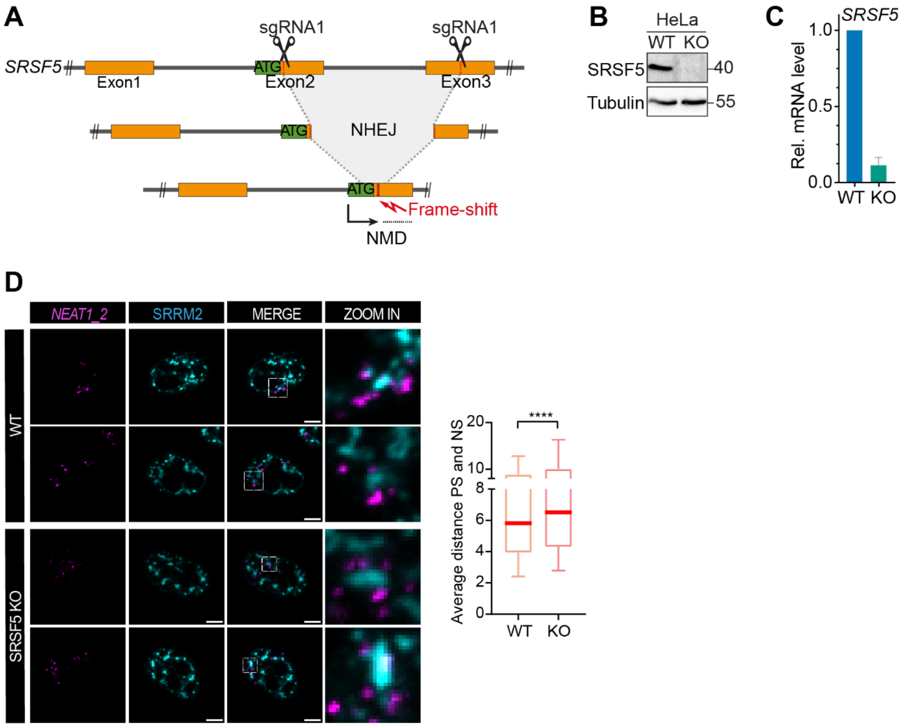


**Figure S3. Long-term depletion or SRSF5 KO triggers a PS compensation mechanism. A)** Generation of a HeLa SRSF5 KO cell line using CRISPR/Cas9. CRISPR genome-editing strategy using two sgRNAs to generate a frame-shift in exon 2 of the *SRSF5* gene and prevent its expression. Nonsense-mediated decay (NMD) degrades the malfunctioning RNA. **B)** Validation of protein KO by Western blot using an SRSF5-specific antibody (a-SRp40). Anti-tubulin probing was used to control for equal loading. **C)** Validation of *SRSF5* mRNA degradation by NMD using RT-qPCR quantifying the *SRSF5* levels relative to U6 snRNA. **D) Left:** Example micrographs of WT and SRSF5 KO cells. PS were labelled with probes hybridizing to the middle region of *NEAT1_2.* NS were detected/immunostained using an anti-SRRM2 antibody. **Right:** Quantification of the distance (pixel) between each PS center of mass to the closest NS center of mass for WT (n = 1275) and SRSF5 KO cells (n = 2055). Mann-Whitney test was performed to evaluate the significance.

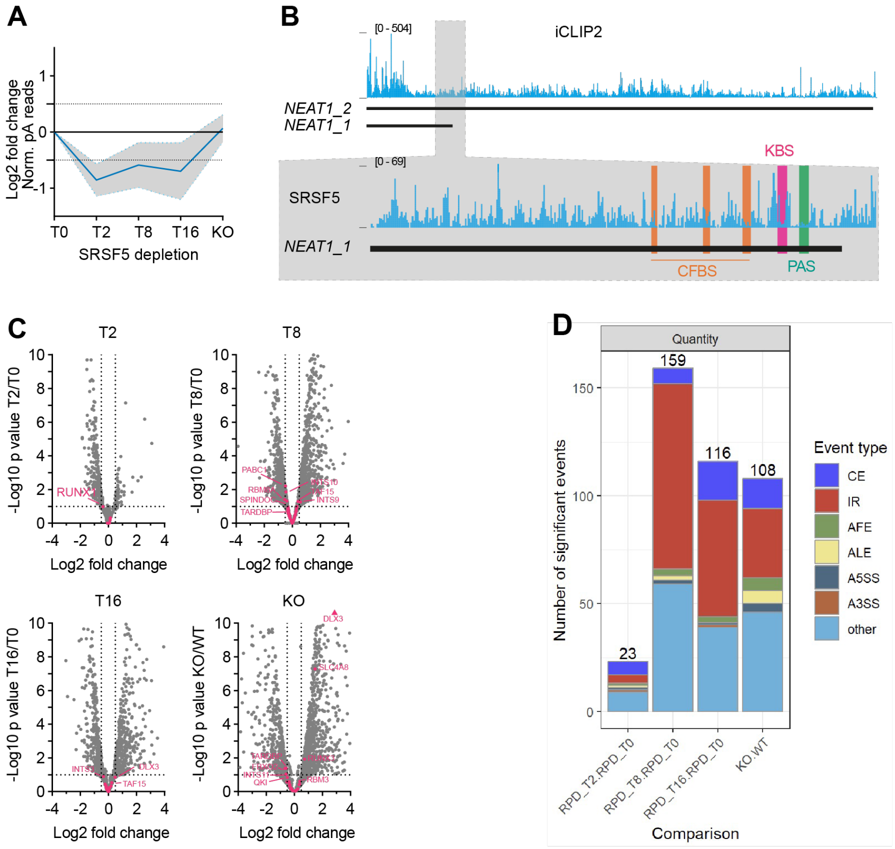


**Figure S4. SRSF5 restricts *NEAT1_2* levels and PS assembly indirectly by regulating TDP-43 and INTS10. A)** Browsershots showing SRSF5 crosslinks on *NEAT1_2* RNA. Zoom-in on the poly(A) site (PAS, green) with known binding sites of cleavage factors (orange, CFBS) or hnRNPK (pink, KBS). **B)** Volcano plots showing differentially expressed genes from the Nascent-seq data at T2, T8 and T16 compared to T0, or SRSF5 KO compared to WT. All genes encoding known PS regulatory factors are indicated. **C)** Splicing changes quantified with MAJIQ from the Nascent-seq data at T2, T8 and T16 compared to T0, or SRSF5 KO compared to WT. CE - cassette exon, IR - intron retention, AFE - alternative first exon, ALE - alternative last exon, A5SS - alternative 5'splice site, A3SS - alternative 3'splice site.

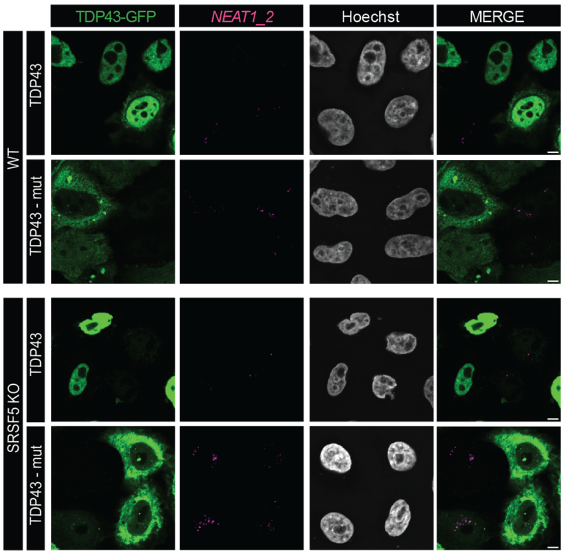


**Figure S5. SRSF5 regulates TDP43 levels.** Representative micrographs showing the subcellular localization of GFP-tagged TDP43 and a mutant TDP43-mut after transfection of HeLa WT and SRSF5 KO cells. TDP43-mut is mutated in the nuclear localization signal of TDP43 (K82A, R83A, K84A, K95A, K96A, R97A). GFP-tagged proteins were expressed from a plasmid. PS were labelled with probes hybridizing to the middle region of *NEAT1_2* and nuclei with Hoechst. Scale bars = 5 μm.

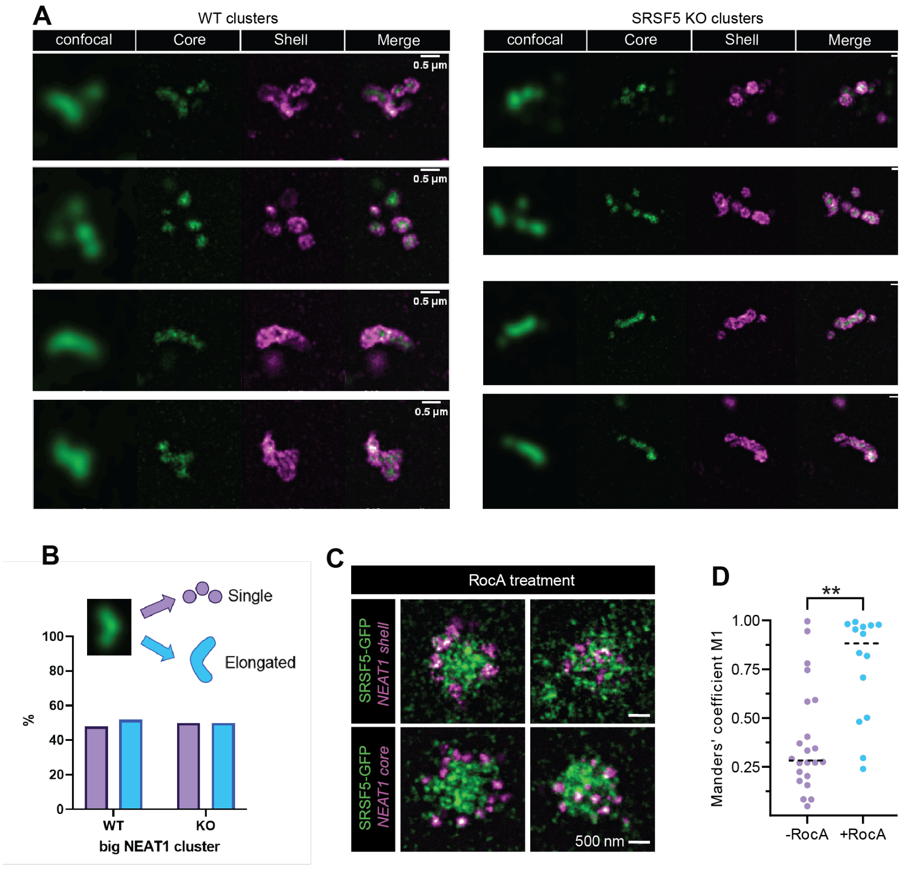


**Figure S6: SRSF5 co-localizes more with PS in hypoxia and RocA-treated cells. A)** Representative micrographs of WT and SRSF5 KO cells showing the different morphologies of PS clusters - elongated rod-like structures or glued PS spheres - using confocal and dSTORM imaging. PS were labelled with probes hybridizing to the middle region (core) or the 5'end (shell) of *NEAT1_2*. Scale bars = 500 nm. **B)** Determination of the percentage of elongated rods or glued PS spheres in WT and SRSF5 KO cells from n = 40 cells. **C)** Cells treated with or without RocA. DNA-PAINT RNA-FISH using probes at the 5'end (shell) or middle region (core) of *NEAT1_2* was coupled with dSTORM imaging for SRSF5-GFP using GFP nanobodies. Scale bars - 500 nm. **D)** Quantification of Manders coefficient indicates complete co-localization of *NEAT1_2* and SRSF5-GFP signal.

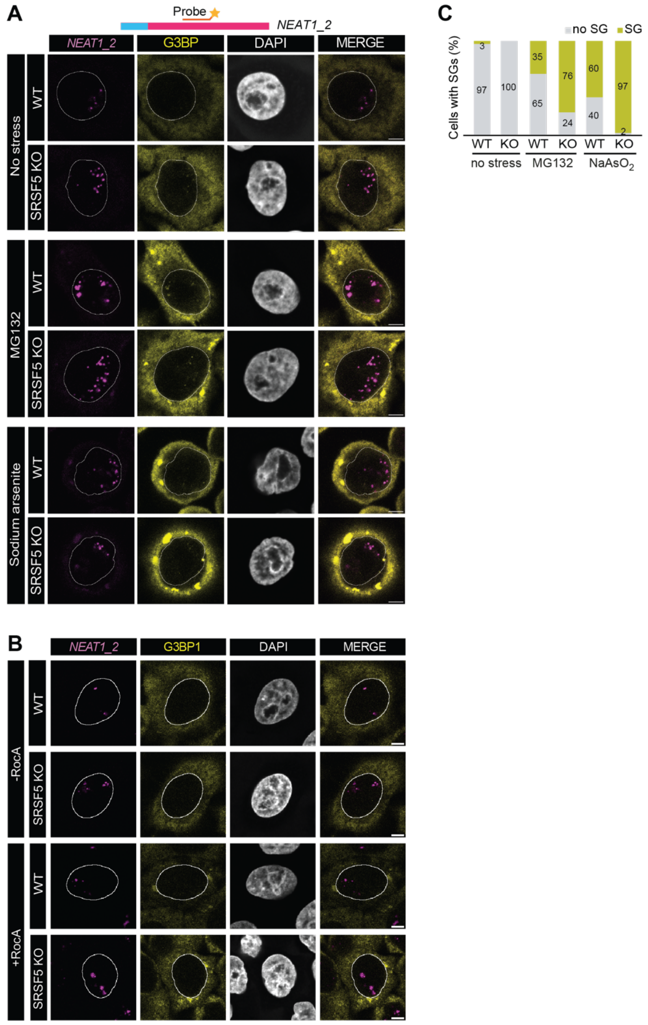


**Figure S7: SRSF5 is required for PS hyper-assembly during stress. A)** Example micrographs of HeLa WT and SRSF5 KO cells subjected to MG132 (4 h, 10 μM) and sodium arsenite (4 h, 0.25 mM). **B)** Example micrographs of HeLa WT and SRSF5 KO cells subjected to RocA (4 h, 5 μM). PS are labelled by RNA-FISH with a probe hybridizing to the middle region of *NEAT1_2*. SGs are labelled by immunofluorescence using an anti-G3BP1 antibody. Nuclei are stained with Hoechst. Scale bars - 5 μm. **C)** Quantification of stress granules SGs.

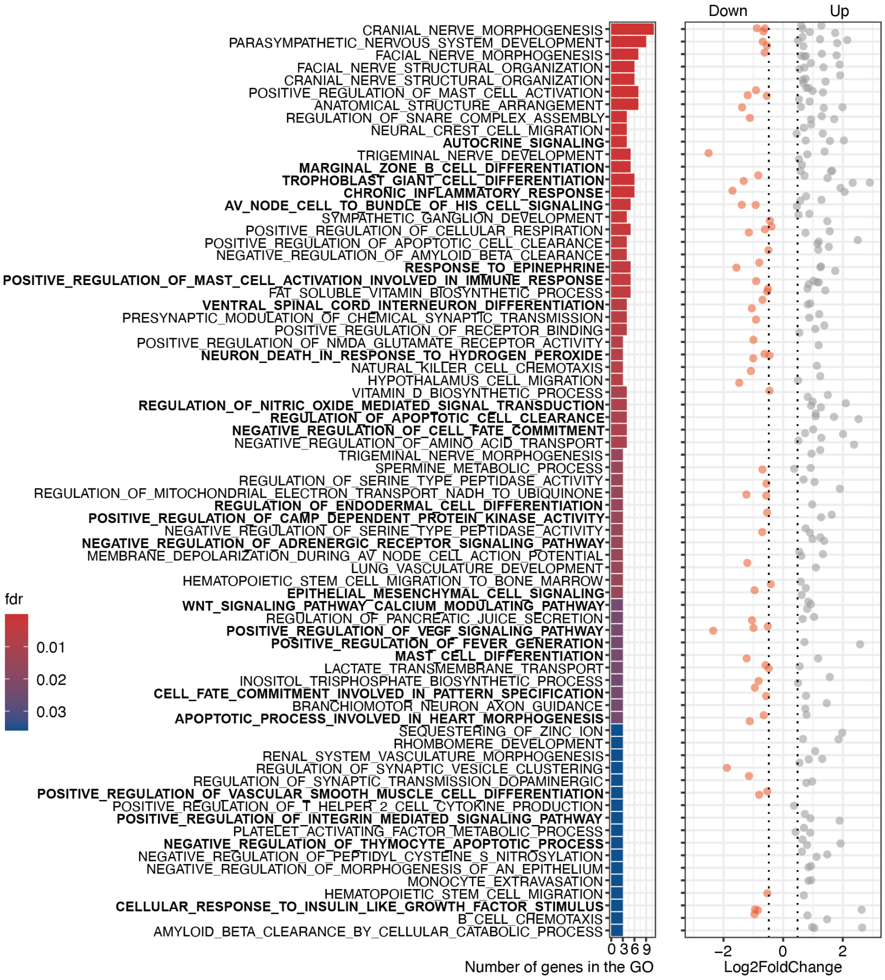


**Figure S8: GO-term analysis of significantly differentially expressed genes in SRSF5 KO cells indicates the dysregulation of many signalling pathways.**

**Supplementary Tables**

**Table S1: List of RNA FISH probes used in this study**

| Target | Fluorophore | Dilution | Supplier | Catalog-Nr. |
|---|---|---|---|---|
| ***NEAT1_2 middle*** | Quasar 670 Dye | 125 nM in Hybridization buffer | BioCat | VSMF-2251-5-BS |
| ***NEAT1_2 5'end*** | Quasar 570 Dye | 125 nM in Hybridization buffer | BioCat | VSMF-3034-5-BS |
| ***NEAT1_2 5'end*** | Quasar 670 Dye | 125 nM in Hybridization buffer | BioCat | VSMF-2246-5-BS |

**Table S2: List of RNA FISH and DNA-PAINT strands for super resolution microscopy**

| Dataset | Name | Sequences 5' to 3' | Modification | Supplier |
|---|---|---|---|---|
| **Figure 3 G** | Imager LP1-AT655 | TAGATGTAT (left handed DNA) | 3'-ATTO 655 | biomers.net |
| | Docking LP1 | TTATACATCTA (left handed DNA) | 5'-Azide | biomers.net |
| **Figure 3 D** | Imager P1-STAR635P | TAGATGTAT | 3'-Abberior STAR635P | biomers.net |
| | Imager P5-STAROrange | ATACATTGA | 3'-Abberior STAROrange | biomers.net |

**Table S3: List of sgRNAs used in this study**

| Species | Name/Target | Protospacer sequence | PAM | Purpose | Supplier |
|---|---|---|---|---|---|
| Human | SRSF5_sg1 crRNA | tactagccggacatcatgag | TGG | NHEJ, KO | IDT |
| Human | SRSF5_sg2 crRNA | gatgctgtgtatgagcttga | TGG | NHEJ, KO | IDT |
| - | universal tracrRNA | - | - | - | IDT |

**Table S4: List of antibodies used in this study**

| Name | Species | Supplier | Catalog-Nr. |
|---|---|---|---|
| α-GFP | Goat | Eric Geertsma, MPI-CBG | - |
| α-INTS10 | Rabbit | Abcam | ab180934 |
| α-SRp40 (SRSF5) | Rabbit | Merck Millipore | 06-1365 |
| α-TDP-43 | Rabbit | Abcam | ab109535 |
| α-alpha-Tubulin | Rabbit | Abcam | ab176560 |
| α-goat-HRP | Donkey | Sigma-Aldrich | AB324P |
| α-rabbit-HRP | Donkey | Merck Millipore | AP182P |
| α-rabbit-Alexa Fluor 680 | Donkey | Thermo Fisher | A10043 |
| α-rabbit-Alexa Fluor Plus 405 | Donkey | Thermo Fisher | A48258 |
| α-rabbit-Alexa Fluor Plus 555 | Goat | ThermoFisher | A32732 |
| α-rabbit-Alexa Fluor Plus 488 | Goat | ThermoFisher | A-11008 |

| | | | |
|---|---|---|---|
| α-mouse-Alexa Fluor Plus 594 | Donkey | ThermoFisher | A-21203 |
| α-rabbit-Alexa Fluor Plus 594 | Donkey | ThermoFisher | A-21207 |
| α-SRRM2 | Rabbit | ThermoFisher | PA5-59559 |

**Table S5: List of Cell lines used in this study**

| Name | Selection | Source | Species |
|---|---|---|---|
| **HeLa wild type** | - | American Type Culture Collection | Human |
| **HeLa SRSF5 KO** | - | David Stanek / This work | Human |
| **HeLa hGRAD NONO-GFP** | Puro/Gen | (Arnold et al., 2024) | Human |
| **HeLa hGRAD SRSF5-GFP** | Puro/Gen | (Arnold et al., 2024) | Human |
| **HeLa hGRAD SRRM2-GFP** | Puro/Gen | (Arnold et al., 2024) | Human |

**Table S6: List of primers used in this study**

| Primer | Primer Sequence (5'-3') | Insert / Amplicon |
|---|---|---|
| qPCR_U6_F | aaatatggaacgcttcacgaatt | *U6* snRNA qPCR Normalization |
| qPCR_U6_R | aggtctagggggaccacagt | *U6* snRNA qPCR Normalization |
| qPCR_hNEAT1_all_F | ctcagctatgcaagagcggc | *NEAT1 all* |
| qPCR_hNEAT1_all_R | tcttctaaagggccgccaca | *NEAT1 all* |
| qPCR_hNEAT1_2_F | gctgaggccagaggaactca | *NEAT1_2* |
| qPCR_hNEAT1_2_R | aggggtttggctttgttcgt | *NEAT1_2* |
| qPCR_TARDBP_F | ggctcatcttggctttgctta | *TARDBP* |
| qPCR_TARDBP_R | tgctgtacgacatgtttgtga | *TARDBP* |
| qPCR_INTS10_F | atttcacactggacccgacc | *INTS10* |
| qPCR_INTS10_R | agtcacgcccttgggg | *INTS10* |
| qPCR_INTS11_F | ccaggtggaagtggctaatga | *INTS11* |
| qPCR_INTS11_R | aaatatggaacgcttcacgaatt | *INTS11* |

**Table S7: List of plasmids used in this study**

| Name | Resistance | Source | Species |
|---|---|---|---|
| **P33_pEGFP-C1_hTDP_Wt-** | Kanamycin | Dorothee Dormann | Human |
| **P38_pEGFP-C1_hTDP-43_83_97AAA_Si65Res** | Kanamycin | Dorothee Dormann | Human |
